# Distinct Acetate Utilization Strategies Differentiate Butyrate and Medium-Chain Carboxylic Acid Producing Chain-Elongating Bacteria

**DOI:** 10.1101/2025.05.02.651941

**Authors:** Ian M. Gois, Connor M. Bowers, Byung-Chul Kim, Robert Flick, Christopher E. Lawson

## Abstract

Chain elongating bacteria (CEB) are a unique guild of anaerobes that upcycle organic waste into valuable short- and medium-chain carboxylic acids (MCCAs), enabling a circular bioeconomy. However, the metabolic rules that determine product chain length have remained elusive. Here, we combine ^13^C isotope tracing, proteomics, enzyme assays, and metabolic modelling to show that distinct acetate utilization strategies underlie the divergence between butyrate- and MCCA-producing CEB. MCCA-producing strains recycle acetate to maximize lactate use under acetate limitation, but at the cost of slower growth. In contrast, butyrate-producing strains grow faster by favoring acetate assimilation, at the cost of restricted lactate utilization when acetate is scarce. These physiological trade-offs are encoded in the substrate specificity of coenzyme A transferase, the terminal enzyme in reverse β-oxidation. Our findings uncover a fundamental constraint shaping chain-length selectivity in CEB and offer new strategies to optimize MCCA production from organic waste streams.

## Introduction

Anaerobic fermentation could transform billions of tons of organic waste per year into economically viable, carbon-negative fuels and chemicals, enabling a circular bioeconomy. One promising approach is microbial chain elongation, which produces C_6_-C_8_ medium chain carboxylic acids (MCCAs)^1^. These versatile platform chemicals can be applied directly in agricultural feeds, nutritional supplements, cosmetics, antimicrobials and surfactants^2,3^, or derivatized to produce liquid fuels^4–6^, polyhydroxyalkanoates^7^, and other oleochemicals such as alcohols, diols and dicarboxylates^8,9^. Because most MCCAs are currently sourced from palm kernel and coconut oil^1^, local production via waste fermentation offers additional environmental benefit by mitigating deforestation and biodiversity loss.

MCCAs are produced by chain elongating bacteria (CEB), a unique guild of strict anaerobes first described by Barker in 1939 with the isolation of *Clostridium kluyveri*^10^. A key challenge in deploying chain elongation industrially has been controlling product distribution. While all known CEB synthesize butyrate (C4), hexanoate (C6) and/or octanoate (C8) via the reverse beta-oxidation (RBO) pathway^11^, the relative abundance of different chain length carboxylates varies drastically among species and environmental conditions. For example, some CEB only produce butyrate, including many abundant in the human gut^12,13^, whereas others produce a substantial amount of hexanoate and octanoate, despite using the same RBO pathway to elongate carbon chains^14–16^. Understanding the ecological and physiological principles that drive some CEB to produce MCCAs rather than only butyrate is therefore essential for improving the yield, selectivity and productivity of MCCA bioprocesses^11^.

During RBO, an electron donor such as lactate is oxidized to acetyl-CoA, which then condenses with an acyl-CoA to initiate a new chain elongation cycle (Figure 1). The resulting intermediate passes through a series of reduction and dehydration steps, elongating the carbon chain by two carbons each cycle. Termination of chain elongation is typically catalyzed by a CoA transferase (CoAT), which transfers CoA from the final acyl-CoA to acetate, yielding a free carboxylic acid (i.e., butyrate, hexanoate, or octanoate) and regenerating acetyl-CoA^17^. Despite a general understanding of RBO, how acetate is used in the pathway and its implications for energy conservation have been debated. Early work by Barker proposed that the acetyl-CoA produced by termination enters the RBO cycle, and energy is conserved via a then-unknown chemiosmotic mechanism. We call this stoichiometry *acetate assimilation*, because it leads to net acetate consumption:

**Figure 1.**
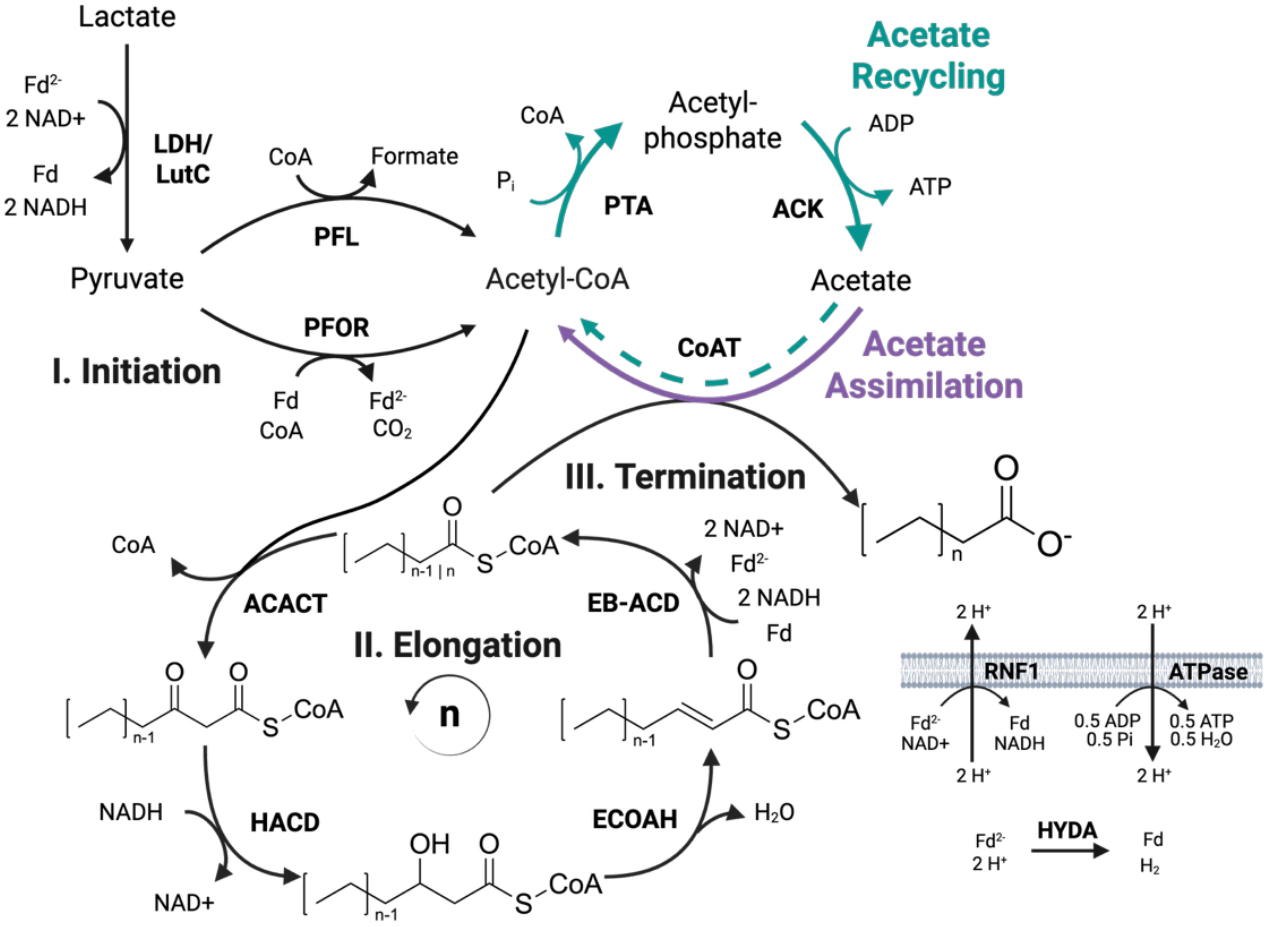
Butyrate (n=1), hexanoate (n=2) and octanoate (n=3) production from lactate. Initiation (I) generates acetyl-CoA via lactate oxidation. Acetyl-CoA is condensed with an acyl-CoA to form a lengthened carbon chain that is reduced and dehydrated during elongation (II) via the RBO cycle. The cycle is terminated (III) by a CoA transferase (CoAT), which transfers CoA from the final acyl-CoA to acetate to form a free carboxylic acid and regenerate acetyl-CoA. Acetyl-CoA may re-enter RBO, resulting in net acetate consumption (acetate assimilation) and energy conservation by chemiosmosis through the RNF complex. Alternatively, acetyl-CoA can be diverted through phosphate acetyltransferase (PTA) and acetate kinase (ACK) to regenerate acetate and conserve energy through substrate level phosphorylation (SLP). Purple arrow indicate reactions leading to acetate assimilation. Green arrows indicate reactions leading to acetate recycling. Biorender.com/wsekjeu.

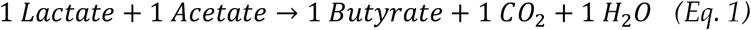

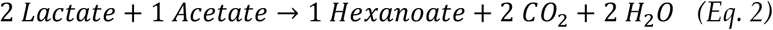

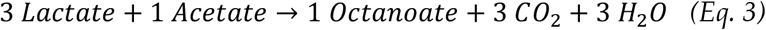

In Barker’s work, the organism under study, *Clostridium kluyveri*, consumed ethanol rather than lactate as its electron donor, but both substrates yield two molecules of NADH when oxidized to acetyl-CoA and therefore follow similar overall stoichiometry. Later, Thauer and colleagues proposed that not all product formation requires net acetate consumption^18–20^. Instead of entering the RBO cycle, some acetyl-CoA is diverted by phosphate acetyltransferase (PTA) and acetate kinase (ACK) to regenerate acetate, while producing ATP via substrate-level phosphorylation (SLP)^11^. We call this stoichiometry *acetate recycling:*

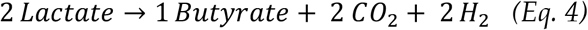

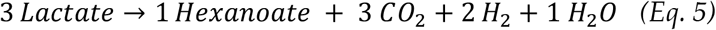

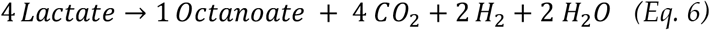

Barker’s hypothesis was later supported–ironically by Thauer *et al*.–with the discovery of chemiosmotic energy conservation in *C. kluyveri*^21–24^. The reduction of an enoyl-CoA is coupled to the reduction of ferredoxin via electron bifurcation, then reduced ferredoxin is oxidized via the membrane bound ferredoxin:NAD^+^ oxidoreductase (RNF complex) which generates ion motive force (IMF) for ATP synthesis by ATPase^21,22^ (Fig. 1). As a result, it is now widely accepted that CEB conserve energy through a combination of both chemiosmosis and substrate-level phosphorylation, with chemiosmosis generally assumed to dominate^2,3,11,21,23,24^. However, this has not been systematically tested across species and growth conditions.

Here, we combine batch fermentations, ^13^C isotope tracing, proteomics, enzyme assays, and metabolic modelling to investigate the relationship between acetate metabolism and product formation in a butyrate-producing CEB (*Anaerostipes caccae*) and a MCCA-producing CEB (*Pseudoramibacter alactolyticus*). Our results suggest that acetate utilization strategy differentiates MCCA-producing and butyrate-producing CEB. We show that MCCA production allows for increased acetate recycling by maintaining redox balance. This enables more lactate utilization under acetate-limiting conditions, at the cost of slower growth due to an increased protein burden associated with producing longer products. In contrast, exclusive butyrate production enables faster growth due to a reduced catabolic protein demand, but constrains acetate recycling, resulting in poor lactate utilization and growth under acetate limiting conditions. This trade-off is reflected by differing substrate chain-length specificities of CoAT in MCCA and butyrate producers. Together, these findings provide foundational insights into chain-length selectivity of CEB and offer new strategies to enhance MCCA production from waste.

## Results

### *P. alactolyticus* maintains growth and lactate consumption during acetate scarcity, producing longer chain-length products

*A. caccae* and *P. alactolyticus* are CEB of great interest. *A. caccae* is an important butyrate producer in the human gut^13,25–27^, and the genus *Pseudoramibacter* is often observed in chain elongation bioreactors producing MCCAs^17,28–33^. To evaluate how acetate availability affects growth, product formation and substrate utilization in *A. caccae and P. alactolyticus*, we conducted lactate batch fermentations with and without acetate. When acetate was supplemented, *A. caccae* only produced butyrate while *P. alactolyticus* produced butyrate and hexanoate, and effectively all lactate was consumed by both organisms. Interestingly, acetate consumption by *P. alactolyticus* was significantly inferior to consumption by *A. caccae*. When lactate was provided and acetate was not supplemented, *A. caccae* did not display significant growth. *P. alactolyticus* grew and fully consumed lactate, albeit at a slower rate than with acetate, and carboxylic acid production shifted towards hexanoate and octanoate (Fig. 2a). Hence, acetate appears to be a mandatory co-substrate for *A. caccae* but a facultative one for *P. alactolyticus*.

**Figure 2.**
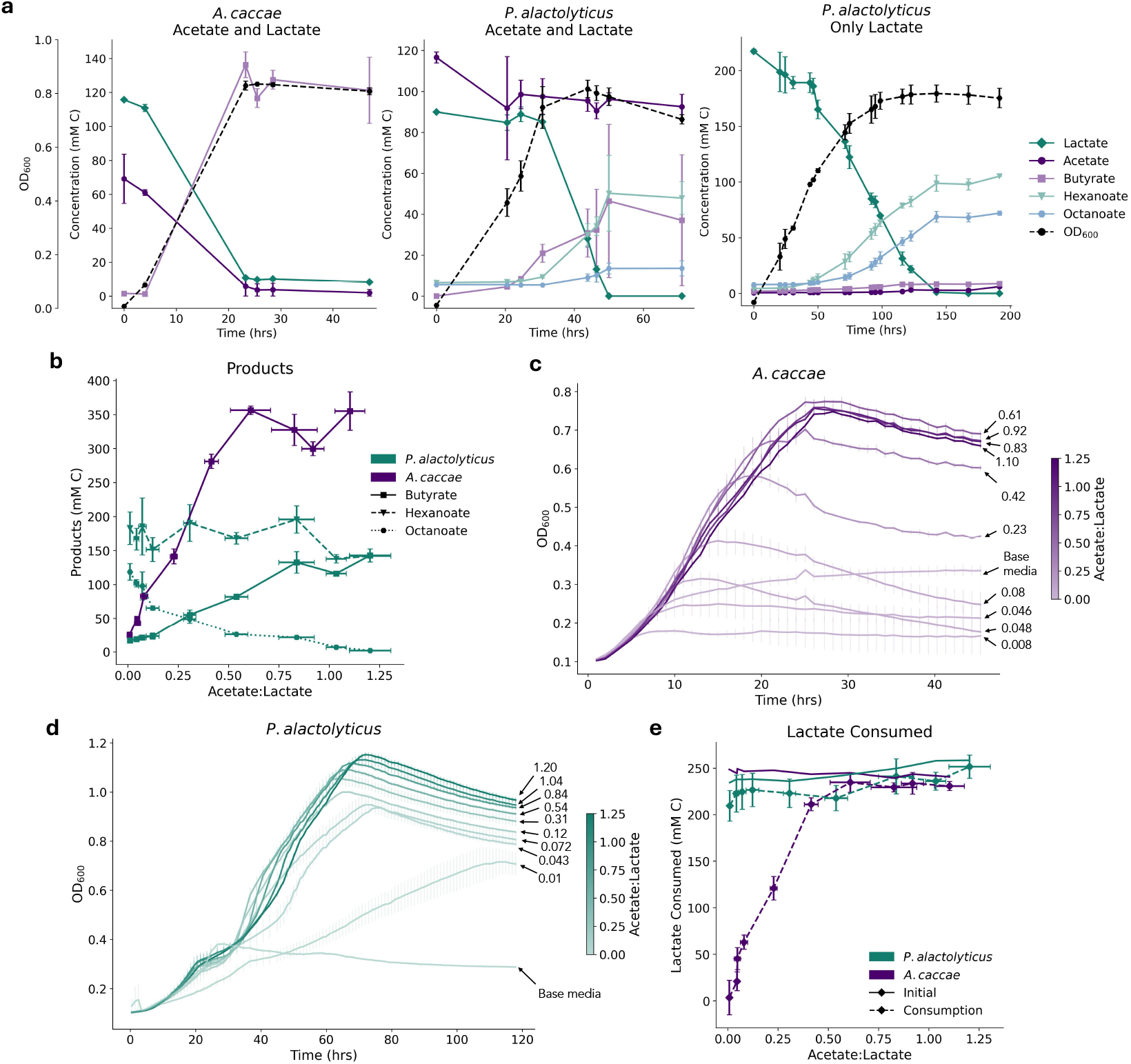
Batch experiments show effects of acetate-to-lactate ratio on carboxylic acid profile, growth and lactate utilization in *A. caccae* and *P. alactolyticus*. **a.** Substrate and product concentrations and growth throughout batch fermentation of *A. caccae* and *P. alactolyticus* on lactate and acetate, and of *P. alactolyticus* on lactate only (n = 3). **b**. Endpoint concentrations of butyrate, hexanoate and octanoate produced by both organisms at different acetate-to-lactate ratios (n = 3). *P. alactolyticus* favours hexanoate and octanoate production as ratio decreases, *A. caccae* solely produces butyrate. Error bars represent 1 standard deviation. **c**. Growth curves of *A. caccae* and **d**. *P. alactolyticus* at varied acetate-to-lactate ratios (lactate concentration fixed at 100 mM) (n = 3). **e**. *P. alactolyticus* maintains growth and lactate consumption at low ratios, whereas these are severely hindered in *A. caccae*. Initial lactate concentrations and lactate consumed (initial minus endpoint concentration) by both organisms at different acetate-to-lactate ratios (n = 3). Base media condition does not contain lactate or acetate supplemented.

We then evaluated differences in these organisms’ responses to intermediate acetate-to-lactate ratios through batch cultures with acetate concentrations varying between 5 mM and 120 mM and a constant lactate concentration of 100 mM. Whereas *A. caccae* only produced butyrate in all conditions, with production increasing with acetate availability, *P. alactolyticus* produced a mixture of butyrate, hexanoate and octanoate, favouring longer products at low acetate-to-lactate ratios (Fig. 2b). Lengthening of products in response to acetate scarcity is reminiscent of similar observations in other MCCA-producing CEB and bioreactors^10,11,34–36^.

*A. caccae* growth decreased significantly as acetate supplementation was reduced (Fig. 2c, Supplementary Material Fig. 1b). On the other hand, *P. alactolyticus* maintained a relatively constant maximum OD as acetate decreased (Fig. 2d, Supplementary Material Fig. 1a). Consistent with these observations, lactate consumption by *A. caccae* was poor at low acetate-to-lactate ratios and did not approach the amount available in the media until a ratio of 0.5, while lactate was effectively fully consumed by *P. alactolyticus* at all tested ratios (Fig. 2e). This confirms that *A. caccae* depends more strongly on acetate than *P. alactolyticus*.

Unexpectedly, we observed minor growth on rich base media without lactate or acetate supplementation (Fig. 2c,d). This is likely due to trace amounts of lactate and acetate in the base media, as well as small amounts of sugars and/or amino acids catabolized by *P. alactolyticus*. However, consumption of background media components by *P. alactolyticus* was negligeable when acetate and lactate were abundant as confirmed by carbon balances (Supplementary Material Fig. 2), indicating that acetate and lactate were the primary substrates for RBO. Interestingly, *A. caccae* growth and lactate consumption was higher on base media compared to media supplemented with 100 mM lactate and 0 to 8 mM (low) acetate (Fig. 2c, Supplementary Material), highlighting this organism’s requirement for acetate as a co-substrate to lactate. To confirm that background media components had minimal effects on the results of our study, experiments were reproduced in dilute media, showing identical trends (Supplementary Material Fig. 3 a,c,e).

**Figure 3.**
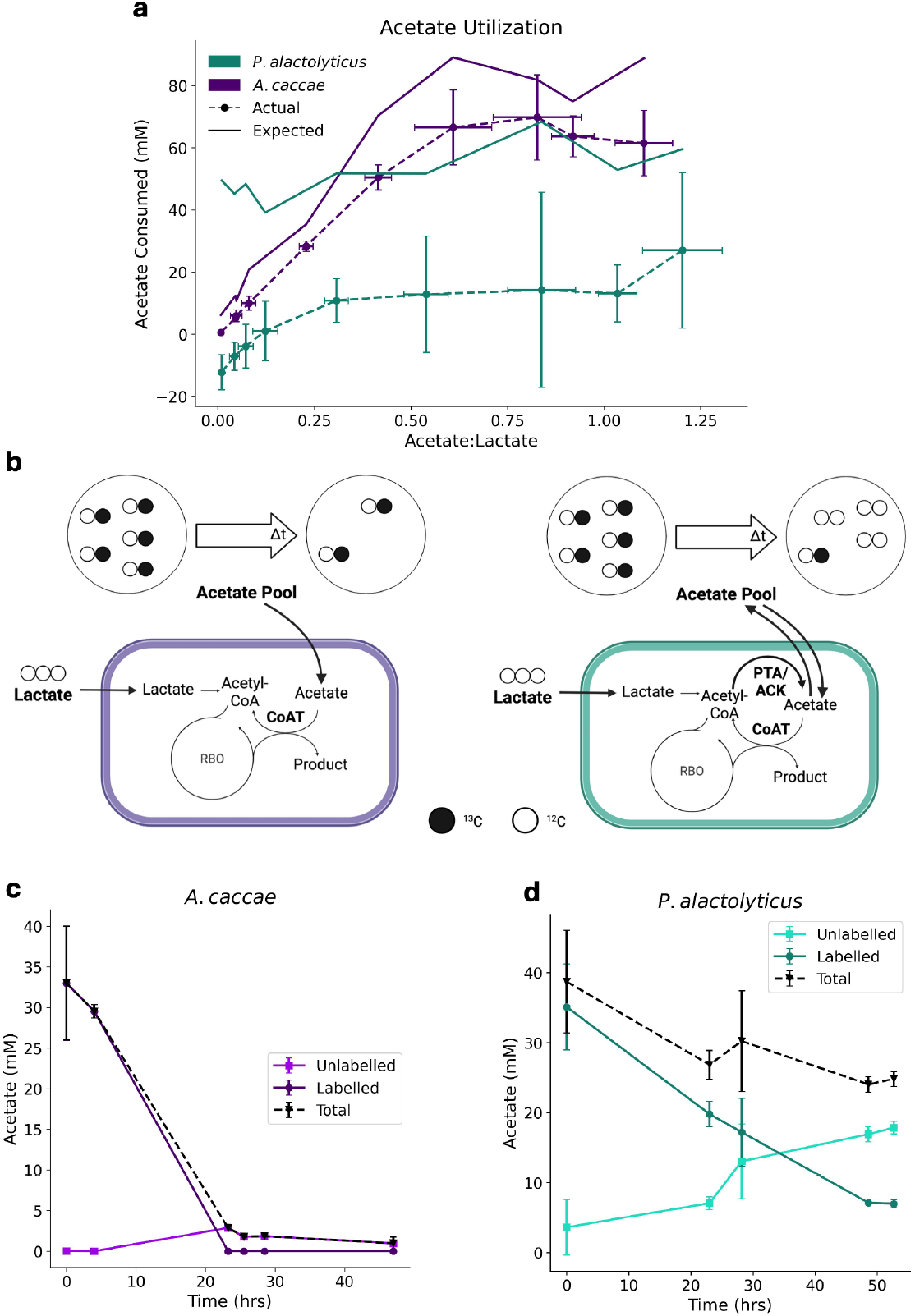
1-^13^C acetate tracing shows different acetate utilization strategies in *A. caccae* and *P. alactolyticus*. **a.** Measured acetate consumption and consumption expected if no recycling were occurring at different acetate-to-lactate ratios (n=3). **b**. Left: Schematics of full acetate assimilation in the tracing experiment. Recycling is minimal and acetate pool labelling pattern does not significantly change after a period of time (Dt). Acetate pool concentration decreases due to assimilation, via CoAT. ^13^C Labeled carbon presented as a black circle. Right: Schematics of a highly active acetate recycling, through acetate regeneration via PTA/ACK. Acetate labelling pattern drastically changes over time, with limited assimilation. Biorender.com/coz8d26. **c**. 1-^13^C-acetate tracing throughout batch growth of *P. alactolyticus*. **d**. and *A. caccae* (n=3). *P. alactolyticus* recycles acetate readily, causing low net consumption, whereas recycling in *A. caccae* is minimal.

### Favouring acetate recycling allows *P. alactolyticus* to maintain growth under acetate scarcity

Acetate recycling through PTA and ACK could allow *P. alactolyticus* to consume less acetate than *A. caccae*, thereby reducing its reliance on it as a co-substrate. To evaluate whether acetate recycling is more prevalent in *P. alactolyticus*, we compared acetate consumption by both organisms across acetate-to-lactate ratios to its expected value if no acetate were recycled. Without recycling, one mole of acetate is consumed for each mole of product formed, so expected acetate consumption is the molar sum of products formed. With *A. caccae*, acetate consumption was slightly inferior to its expected value for all tested ratios, suggesting minimal recycling (Fig. 3a). On the other hand, acetate consumption by *P. alactolyticus* was significantly lower than expected if one mole of acetate was assimilated for each mole of product. In fact, minimal production was detected at low ratios (Fig. 3a), indicating that acetate recycling by *P. alactolyticus* is superior to that by *A. caccae*.

To confirm this, we performed a tracing experiment by supplementing 1-^13^C labeled acetate to *A. caccae* and *P. alactolyticus* during lactate fermentation (Fig. 3b). In this experiment, we anticipated two possible outcomes. In the first, consumed acetate would be assimilated into carboxylic acids, with any remaining acetate being labeled. This would be indicative of minimal to no recycling. In the second, labeled acetate would be incorporated into carboxylic acids while unlabeled acetate would be produced via unlabeled lactate oxidation and PTA/ACK reactions. Then, labeled acetyl-CoA pool would be substantially diluted, resulting in an extracellular acetate pool containing both unlabeled and labeled acetate. This would indicate strong acetate recycling. The first scenario was observed in *A. caccae* (Fig. 3c), where labeled acetate was depleted and an insignificant amount of unlabeled acetate was generated, indicating minimal but detectable recycling. We observed the second scenario in *P. alactolyticus* (Fig. 3d), where most of the 1-^13^C-acetate was assimilated and the major fraction of the extracellular acetate pool was unlabeled. A similar result has been obtained with *C. kluyveri* using labeled ethanol and acetate^37^. These results support that *P. alactolyticus*, unlike *A. caccae*, does not depend strongly on acetate as a co-substrate because acetate is readily recycled through PTA/ACK, reducing net consumption requirements.

### Favouring acetate recycling requires long chain products for redox balance and reduces chemiosmosis

We then asked whether either organism’s bias towards acetate assimilation or recycling shifted across acetate-to-lactate ratios. Also, it remained unclear why *A. caccae* was incapable of strongly recycling acetate, and how acetate utilization affected product spectra and energy conservation. To address these questions, we used a stoichiometric model of RBO (Fig. 4a, Stoichiometric Model in Methods) to quantify moles of acetate recycled per mole of product (*θ*). This established that an increase in RBO cycles (*n*) is required to recycle acetate strongly (*θ* > 0.5). Model fluxes were calculated based on substrate consumption and product formation measurements across a wide range of acetate-to-lactate ratios (0.08 – 1.2 mol/mol in base ATCC 2107 media, Fig. 2b,e, Fig. 3a; 0.02-1 mol/mol in dilute ATCC2107 media, Supplementary Material Fig. 4).

**Figure 4.**
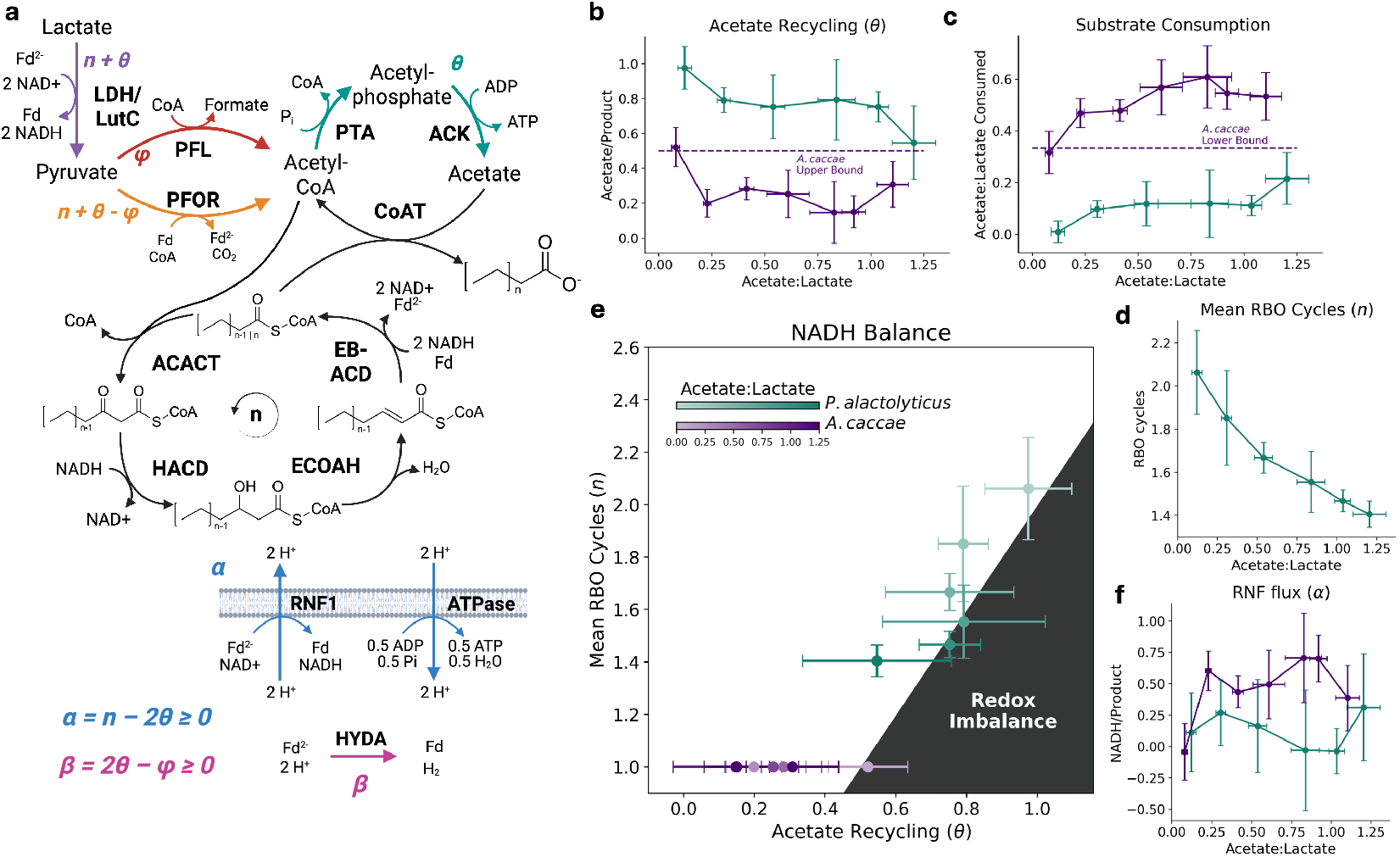
Stronger recycling requires more RBO cycles on average and reduces chemiosmosis. **a.** Stoichiometric model of RBO with simultaneous acetate assimilation and recycling, where θ is the extent of acetate recycling (moles recycled per mole product) and *n* is the mean number of RBO cycles. LDH = electron bifurcating lactate dehydrogenase; LutC = lactate utilization protein C; PFOR = pyruvate:ferredoxin oxidoreductase; PFL = pyruvate formate lyase; PTA = phosphate acetyltransferase; ACK = acetate kinase; ACACT = acetyl-CoA C-acetyltransferase; HACD = 3-hydroxyacyl-CoA dehydrogenase; ECOAH = enoyl-CoA hydratase; EB-ACD = electron bifurcating acyl-CoA dehydrogenase, comprising acyl-CoA dehydrogenase (ACD) and electron transfer flavoproteins A and B (EtfA and EtfB); CoAT = acyl-CoA:acetate CoA transferase; Fd = ferredoxin; RNF1 = proton translocating ferredoxin: NAD+ oxidoreductase complex; ATPase: F0/F1 ATP synthase; HYDA = Fe–Fe hydrogenase. BioRender.com/r3otw9j. **b**. Acetate recycling as a function of acetate-to-lactate ratio for *P. alactolyticus* and *A*. caccae. **c**. Ratio of acetate-to-lactate consumption by *P. alactolyticus* and *A. caccae* across ratios of supplemented acetate and lactate. **d**. Mean number of RBO cycles (n) in *P. alactolyticus* as a function of acetate-to-lactate ratio. **e**. Experimentally derived pairs of n and θ, and inequality relating these parameters for maintenance of redox balance (α > 0). When acetate is scarce, production of longer-chain products by *P. alactolyticus* enables strong acetate recycling (θ > 0.5) by balancing the reduction equivalents released by increased lactate oxidation, thereby maintaining null or positive flux through RNF. Since *A. caccae* only produces butyrate, redox balance is not possible for θ > 0.5, meaning acetate and lactate must be consumed in a ratio of at least 1/3. **f**. Flux through RNF relative to product flux (α).

When acetate was plentiful, acetate recycling by *A. caccae* was minimal (*θ* ≈ 0.2). In this case, NADH produced from lactate is inferior to NADH consumed by the single RBO cycle which synthesizes butyrate (*n* = 1). The reduced ferredoxin produced by electron bifurcation in RBO drives the RNF complex to regenerate and balance NADH. Acetate recycling by *A. caccae* increased when less acetate was supplemented (Fig. 4b). This facilitated lactate utilization, since acetate recycling reduces the ratio in which acetate and lactate are consumed (Fig. 4c). When 0.5 moles of acetate are recycled for each mole of butyrate produced (*θ* = 0.5), NADH production from lactate matches consumption by RBO and RNF flux goes to 0. Assuming that reverse flux through RNF is infeasible, we hypothesized that *A. caccae* could not achieve *θ* > 0.5 because NADH would accumulate. This means *A. caccae* would need to consume acetate and lactate in a ratio of at least 1/3. Indeed, *θ* was smaller or equal to 0.5 and *α* was larger or equal to 0 in all conditions, and the ratio of acetate-to-lactate consumption did not fall significantly below 1/3. This lower bound explains why growth and lactate utilization by *A. caccae* dropped sharply between acetate-to-lactate supplementation ratios of 0.5 and 0.25 (Fig. 2c,d).

On the other hand, *P. alactolyticus* maintained growth and full lactate utilization when acetate was scarce. As suggested previously, this is due to its enhanced capacity for acetate recycling: *P. alactolyticus* achieved *θ* ≥ 0.5 in all tested conditions, with full recycling (*θ* ≈ 1) occurring at low acetate-to-lactate ratios. This was impossible in *A. caccae* as NADH from lactate oxidation would quickly accumulate, but unlike *A. caccae, P. alactolyticus* can produce hexanoate and octanoate in addition to butyrate. As these MCCAs require more NADH, stronger acetate recycling is possible without the loss of redox balance. Indeed, *P. alactolyticus* favoured MCCA production as acetate supplementation was reduced and *θ* approached 1 (Fig. 4b,d,e). Heightened NADH production from lactate was counteracted by increasingly lengthened products (increasing *n*) and predicted RNF flux remained within a 95% CI of 0 (Fig. 4f). Thus, we propose that by favouring MCCA production when acetate is scarce, *P. alactolyticus* can fully recycle acetate while maintaining redox balance. This explains why, unlike *A. caccae, P. alactolyticus* does not require acetate as a co-substrate and maintains full lactate utilization when acetate is scarce. Increased acetate recycling accompanied by an increase in product chain length was also observed in two other CEB, *Clostridium kluyveri* and *Megasphaera hexanoica*, when we grew them in diluted ATCC 2107 medium (Supplementary Material Fig. 4). Interestingly, *C. kluyveri* recycled acetate less than *P. alactolyticus* in acetate limited conditions (*θ* of 0.60 and 1.21, respectively) (Supplementary Material Fig. 4b, Supplementary Material S1). *C. kluyveri* also ran fewer RBO cycles than *P. alactolyticus* (*n* of 1.82 and 2.26, respectively) (Supplementary Material Fig. 4d, Supplementary Material S1), resulting in negligeable octanoate production. This is consistent with previous findings with *C. kluyveri*^38,39^. On the other hand, *M. hexanoica*, like *P. alactolyticus*, favoured octanoate production as acetate recycling increased (Supplementary Material Fig. 3f). These results indicate that octanoate production enhances capacity for acetate recycling.

Acetate recycling may also increase hydrogen production (Supplementary Material Fig. 5). As was established, acetate recycling enables more lactate utilization when acetate is limiting. Stoichiometrically, this releases more NADH, reducing flux through the RNF complex via product inhibition. The resulting accumulation of reduced ferredoxin can be balanced with hydrogen production by HYDA flux (*β*).

**Figure 5.**
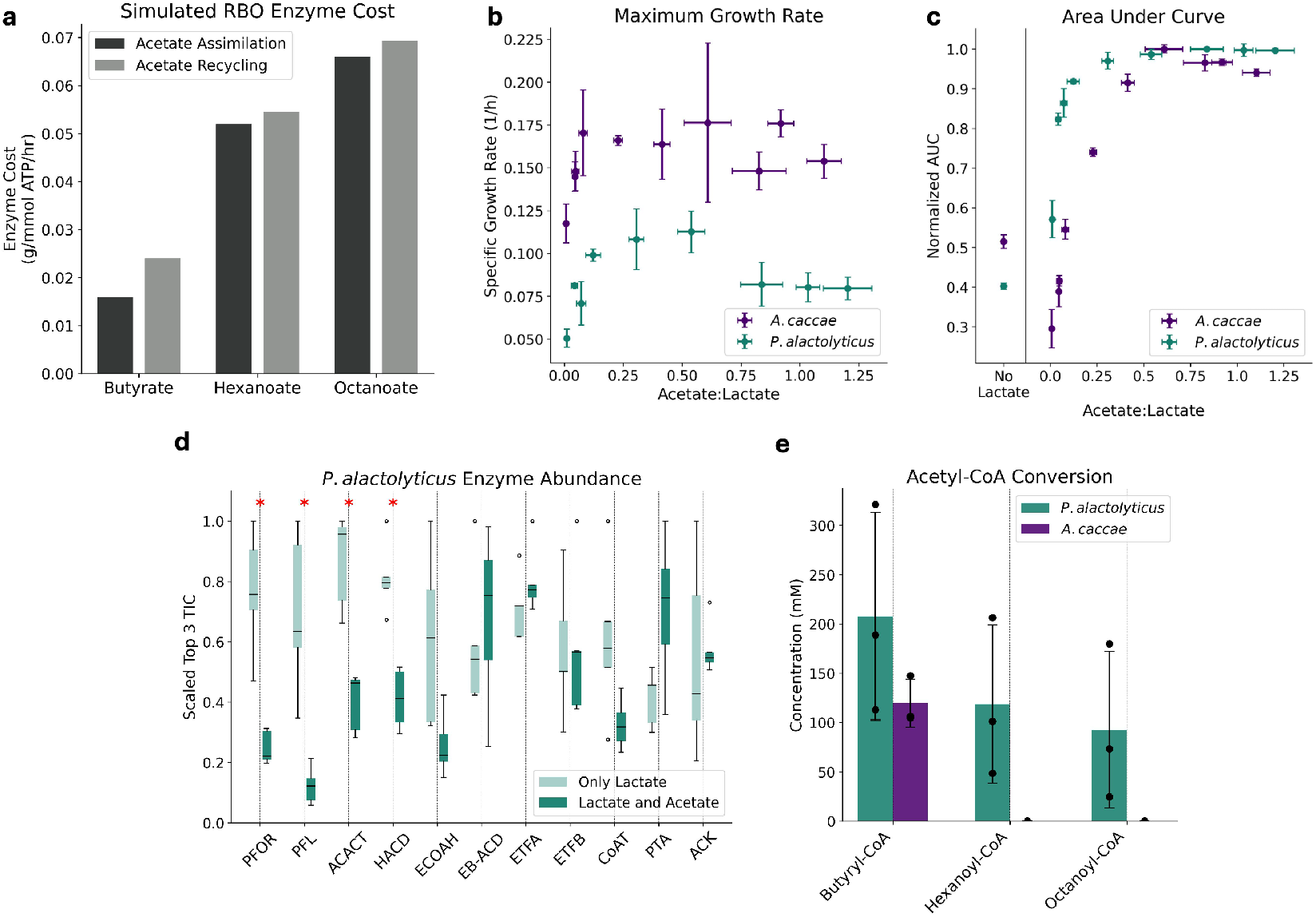
Trade-off between efficient acetate utilization and metabolic burden/growth rate and the restriction of *A. caccae*’s CoAT to acyl-CoA beyond C4. **a.** Predicted enzyme cost per unit ATP flux (g/mmol ATP/hr) required to produce butyrate, hexanoate and octanoate with and without acetate recycling **b**. Maximum growth rates achieved by *P. alactolyticus* and *A. caccae* at different acetate-to-lactate ratios. **c**. Area under the growth curves for *P. alactolyticus* and *A. caccae* **d**. Differential proteomics on *P. alactolyticus* grown on lactate with and without acetate (n=5). Top 3 Total Ion Current (Top 3 TIC) was used as comparison metrics. Each data point was scaled by the maximum value of the corresponding enzyme across all conditions.* p < 0.05. **e**. Acyl-CoA CoA transfer reaction using *A. caccae*’s and *P. alactolyticus*’ cell lysate. All replicates are biological. Recycling acetate and/or producing longer products requires a larger pool of RBO enzymes, likely contributing to the growth rate penalty assumed by *P. alactolyticus* as compared to *A. caccae* and by both organisms when acetate-to-lactate ratio is reduced. *P. alactolyticus*’ CoAT reacts with C4-C8 acyl-CoAs, while *A. caccae*’s is limited to C4.

Alternatively, reduced ferredoxin production can be tempered by redirecting pyruvate from PFOR to PFL: when forming acetyl-CoA, PFOR reduces ferredoxin and produces CO_2_, whereas PFL only produces formate. The puzzling redundancy of PFL and PFOR may then be functional. Flux through PFL (*φ*) allows for some electrons to be balanced with formate rather than hydrogen when acetate is recycled, potentially preventing rapid accumulation of H_2_. Heightened hydrogen partial pressure has been shown to hinder chain elongation due to thermodynamic constraints^40^. Notably, if flux through HYDA must be positive or null, formate production requires some acetate recycling. Otherwise, there would not be enough reduced ferredoxin to regenerate NADH for RBO through RNF. Indeed, formate measurements from select acetate-to-lactate ratios predict non-negative HYDA flux for both organisms (Supplementary Material Fig. 6). PFL flux is higher in *P. alactolyticus* than *A. caccae*, particularly at low acetate-to-lactate ratios when acetate recycling is more prevalent and more ferredoxin must be balanced.

It has been reported that in other chain elongators such as *C. kluyveri*, the HACD enzyme in RBO can use NADPH instead of NADH to reduce oxoacyl-CoAs. Here, the NFN complex would regenerate NADPH from NADH and reduced ferredoxin through electron confurcation^22,41^. This would increase the number of RBO cycles required for acetate recycling, since less NADH is balanced by each RBO cycle (see stoichiometric model in *Methods*). Our data suggests that NFN does not play a major role in RBO for *P. alactolyticus* and *A. caccae*, since these organisms satisfy the redox constraint predicted when NFN flux is negligeable (Supplementary Material Fig. 7 for further discussion). However, NFN was found in both proteomes (Supplementary Material S1) and likely plays a role in generating NADPH for anabolism.

Acetate recycling also has important implications for energy conservation in RBO. Chemiosmosis is often assumed to be the primary mode of energy conservation in CEB, with SLP playing a minor and static role. These results show that SLP becomes increasingly prevalent in both *A. caccae* and *P. alactolyticus* as acetate supplementation is reduced. Notably, stoichiometric analysis suggests that acetate recycling reduces RNF flux and ATP yield on lactate (see Stoichiometric Model in Methods), with predicted RNF flux and ATP yield being lower at low acetate to lactate ratios for both *P. alactolyticus* and *A. caccae* (Fig. 1f, Supplementary Text). Taken together, these results suggest that acetate recycling imposes an ATP yield penalty and can require MCCA production (for *θ* > 0.5). Under acetate limitation, increased lactate utilization enabled by recycling may outweigh these potential drawbacks.

### Acetate recycling and requisite production of longer chain products impose higher metabolic burden, reducing growth rate

In addition to energy yield, a critical characteristic of a metabolic strategy is enzyme cost^42–44^. Each reaction in a metabolic pathway demands the synthesis and maintenance of an enzyme pool. Given an arbitrary pathway flux, the size of this pool is a function of the enzyme’s saturation and the reaction’s thermodynamic driving force. It follows that a longer pathway and/or a pathway with more reactions near equilibrium will require more enzyme per unit flux. Resource allocation arguments suggest that a higher catabolic enzyme cost per ATP flux is likely to slow growth, since fewer resources (i.e., biomass) are free to engage in anabolism^8^.

Enzyme Cost Minimization (ECM)^43^ was used to approximate a lower bound for RBO enzyme cost per ATP flux for butyrate, hexanoate or octanoate production with full acetate recycling or full assimilation. ECM predicted that thiolase (ACACT) reactions are the main thermodynamic bottleneck in RBO (Supplementary Material Fig. 8). At the total enzyme cost minimum, the effects of this bottleneck are spread across other reactions, including HACD, ECOAH and EB-ACD, increasing their enzyme cost. Thiolase reactions are increasingly unfavourable as chain length grows (Supplementary Material Fig. 8), therefore MCCA production engenders a higher total enzyme cost than butyrate production. Acetate recycling further heightens enzyme cost due to PTA and ACK and reduced ATP yield on lactate (Fig. 5a, Supplementary Material Fig. 9). Therefore, acetate recycling likely imposes a growth rate penalty, particularly if MCCAs are produced to strongly recycle acetate (*θ* > 0.5). This is consistent with the faster specific growth rates of *A. caccae* vs. *P. alactolyticus* measured from batch cultures with different acetate to lactate ratios (Fig. 2b,c, Fig. 5b): *A. caccae* minimizes enzyme cost by favouring acetate assimilation and only producing butyrate, while *P. alactolyticus* assumes the higher enzymatic burden required produce MCCAs, enabling strong acetate recycling. This allows *P. alactolyticus* to maintain growth and lactate consumption when acetate is scarce (Fig. 5c, Fig. 2d).

Since ECM predicts enzyme cost per unit flux, a higher cost could be reflected in both heightened protein expression and/or reduced flux. To test whether favouring acetate recycling and MCCA production is facilitated by changes in protein expression, we performed differential proteomics analysis on *P. alactolyticus* (Fig. 5d) with and without acetate supplementation. We observed increased expression in PFOR and PFL when acetate was not provided. This may facilitate more pyruvate flux to acetyl-CoA, compensating for increased diversion of acetyl-CoA to acetate recycling. Additionally, we observed heightened expression of ACACT and HACD when acetate was not supplemented and MCCA production was favoured, consistent with ACACT reactions being more thermodynamically constrained for MCCAs. While ECM suggested that MCCA production imparts higher demand for ECOAH and EBACD, expression of these enzymes did not change significantly. This discrepancy is not unexpected, since ECM finds an optimal distribution of metabolite concentrations that may not reflect intracellular conditions. These enzymes may also be more catalytically efficient than expected. Similarly, expression of CoAT, PTA and ACK did not change significantly. Nonetheless, these results support that favouring MCCA production when acetate is scarce imposes an enzyme cost penalty and involves gene expression control in *P. alactolyticus*.

### *A. caccae*’s CoAT activity is limited to butyryl-CoA, preventing further elongation to MCCAs

Despite the impact of acetate utilization strategies on product chain length, lactate utilization, energy conservation, growth and gene expression, the molecular mechanism that constrains *A. caccae*’s products to butyrate while enabling *P. alactolyticus* to synthesize MCCAs remains unclear. One possible explanation could be enzyme specificity. In the synthetic RBO in *E. coli*, thioesterases are employed to terminate elongation cycles by acting on CoA/ACP intermediates, and they are crucial for controlling carboxylic acid chain length^9,45,46^. Analogously, CoA transferases terminate elongation cycles in CEB and may have a similar role in specifying chain length. Some CoATs have been heterologously expressed and characterized in *E. coli*, where it was observed that CoATs from butyrate-producing CEB have preference for butyryl-CoA and no activity on hexanoyl-CoA^47–49^.

To investigate CoAT specificity in *A. caccae* and *P. alactolyticus*, we obtained cell lysates from both organisms and provided acetate and butyryl-CoA, hexanoyl-CoA or octanoyl-CoA as substrates. With *A. caccae*’s lysate, activity was observed on butyryl-CoA but not hexanoyl-CoA. *P. alactolyticus* lysate, on the other hand, exhibited activity not only on butyryl-CoA but also hexanoyl-CoA and octanoyl-CoA (Fig. 5e). Notably, our proteomics data indicate that a single allele is responsible for CoAT activity in both *A. caccae* and *P. alactolyticus*, although multiple CoAT and thioesterase genes are encoded by both organisms. Nonetheless, these results suggest that CoAT specificity is strict in *A. caccae* and broader in *P. alactolyticus*, and providing the first report of CoAT activity on octanoyl-CoA.

## Discussion

While the role of acetate in CEB was debated in the decades following their discovery in 1939^10,18,37,40,50–52^, the consensus has since become that acetate is mainly assimilated, with IMF being the primary mode of energy conservation. Our results challenge this view, as we show that while the butyrate-producer *A. caccae* favours acetate assimilation and energy conservation via IMF, *P. alactolyticus* favours acetate recycling and SLP, producing MCCAs to balance additional redox equivalents from increased lactate oxidation. As a result, *A. caccae* must consume acetate as a co-substrate, whereas *P. alactolyticus* can grow and consume lactate without net acetate consumption. While the present work has focused on lactate-based chain elongation, the stoichiometry of redox equivalents for ethanol-based chain elongation is highly analogous—in both cases, 2 moles of NADH are produced per mole of electron donor. Consequently, ethanol-consuming CEB such as *C. kluyveri* also produce MCCAs to recycle acetate when it is scarce, as evidenced by supplementary experiments (Supplementary Material Fig. 4b-4d) and in previous open culture studies^53,54^. Specific redox balances with other electron donors used by CEB (sugars^36,55^, methanol^56,57^, propanol^58^) will vary. However, in all cases recycling acetate will increase electron donor oxidation. Therefore, redox balance under acetate recycling is likely to require synthesis of longer-chain, more reduced products.

We show that the specific CoAT enzyme variants encoded by CEB are critical for controlling sole butyrate versus MCCA production. In *A. caccae*, CoAT is specific to butyryl-CoA as has been previously shown for other CEB^47,48^, whereas in *P. alactolyticus* it acts on butyryl-CoA, hexanoyl-CoA and octanoyl-CoA (Fig. 5e). In addition to CoAT, ACACTs (thiolases) from *M. hexanoica* and *C. kluyveri* have been shown to enhance C6 production in *E. coli*, suggesting that these enzymes also present chain-length specificity^59,60^. Conventional top-down microbiome engineering approaches that rely on selection pressure and manipulation of bioprocess parameters do not consider how these genetic differences constrain process performance. Using CoAT gene sequences as biomarkers for MCCA-production potential could track MCCA-producing CEB in bioreactors and improve inoculum selection. Moreover, engineering these enzymes for greater control of MCCA chain length will be a key molecular intervention to boost product selectivity.

Substrate specificity of CoAT helps explain why *P. alactolyticus* can synthesize MCCAs and *A. caccae* cannot, affording the former enhanced capacity for acetate recycling and efficient lactate utilization. However, CoAT specificity does not explain how acetate recycling is increased in response to acetate limitation. Favouring acetate recycling may be a passive process, where the reduction of intracellular acetate concentration during acetate limitation increases the thermodynamic driving force and net flux through PTA and ACK compared to lactate oxidation. Likewise, CoAT specificity in *P. alactolyticus* is likely not responsible for favouring MCCA production when acetate is limiting, since proteomic data suggests that *P. alactolyticus* uses a single CoAT for C4, C6 and C8 carbon intermediates (Fig. 5d). Instead, increased MCCA production appears to involve gene regulation to modulate RBO enzyme levels. Through differential proteomic analysis, we observed that ACACT and HACD abundances increased significantly in *P. alactolyticus* when acetate was not supplemented and MCCA production increased. This is consistent with ACACT being the main thermodynamic bottleneck in RBO^61^.

The mechanism for upregulating RBO enzyme expression likely involves the redox-sensing transcriptional regulator Rex, which responds to changes in NADH/NAD^+^ ratios. If acetate recycling increases passively in response to acetate limitation, there would likely be a transient excess of NADH which triggers Rex. The Rex consensus binding site^62,63^ has been identified in the promoter regions of RBO enzyme genes, including ACACT and HACD, in *C. kluyveri*^62^, *Caproicibacterium lactatifermentans*^36^ and *Clostridium acetobutylicum*^63^. In *P. alactolyticus* genome, sequences nearly identical to the consensus can be found upstream of the *acact-ecoah-hacd* operon and the *coat* gene (Supplementary Material S1), suggesting that Rex has an important role in the regulation of RBO genes. As ACACT (initiation) and CoAT (termination) compete for the same substrate, the increase in ACACT and HACD highlighted in our differential proteomics would favor elongation over termination, resulting in longer products. Alternatively, acetylphosphate and acetyl-CoA could be involved in this regulation, as their intracellular concentrations would decrease in response to lower acetate availability. Acetylphosphate is a known signalling molecule in two-component systems^64–66^, while both acetylphosphate and acetyl-CoA participate in lysine acetylation, a post-translational modification^67,68^.

Aside from underlying regulatory mechanisms and enzymatic specificities, capacity for acetate recycling may represent a fundamental ecological distinction between butyrate-producing and MCCA-producing CEB. We suggest that, since butyrate producing CEB cannot strongly recycle acetate, their growth and lactate consumption is limited by acetate availability, as shown above with *A. caccae* (Fig. 1d). On the other hand, MCCA producers are capable of strong acetate recycling. This allows them to maintain lactate utilization when acetate is scarce, demonstrated here with *P. alactolyticus* (Fig. 1d). As a result, MCCA-producing CEB may outcompete butyrate-producing CEB when acetate concentration or feeding rate is low. When acetate is abundant, butyrate production would likely outcompete MCCA production, since butyrate production requires less enzyme investment and may therefore enable faster growth. Accordingly, we propose that environments with low acetate-to-lactate ratios select for MCCA producing CEB, whereas those with high acetate-to-lactate ratios select for butyrate producing CEB.

The environments from which *P. alactolyticus* and *A. caccae* were isolated support this hypothesis. *P. alactolyticus* was isolated from human infection sites^69^ where lactate is likely to be enriched, such as infections of the lungs^70^, brain^71^, and teeth^72^. Conversely, *A. caccae* was isolated from the human gut where acetate accumulates to high concentrations (>50 mM) and is the most abundant SCCA^73–76^. More generally, lower acetate-to-electron donor supplementation ratios promoted MCCA production over butyrate in open culture batch reactors, although propionate production appears to have tempered MCCA production at very low ratios^34,77,78^. Lower ratios of acetate-to-lactate feed rates favoured hexanoate production over butyrate in a continuously fed open culture reactor^78^, consistent with MCCA production enabling increased acetate recycling for faster lactate utilization when acetate feed rate is limiting. Contrary to our expectation, in a semi-continuous chain elongation reactor fed lactate and xylan at constant rates without acetate, butyrate producers eventually outcompeted MCCA producers. However, this shift was associated with the enrichment of potential acetate producers (*Syntrophococcus*), which could have increased acetate cross-feeding and given a competitive advantage to butyrate producers^79^. Overall, these bioprocess studies support that acetate utilization strategy is a key ecological differentiator of butyrate and MCCA-producing CEB, with the latter’s capacity for acetate recycling being advantageous when acetate is limiting.

Our findings point to key genetic, physiological and ecological determinants of MCCA production, supporting new bioengineering and bioprocess developments. Key limitations of conventional MCCA production with microbiomes are the lack of control over microbial interactions and genetics, causing poor MCCA yield and selectivity. Our work suggests that tailoring microbiome function to limit acetate cross-feeding would favour MCCA production over butyrate. We also show that CoAT enzymes from CEB have evolved different substrate preferences, which creates an opportunity to rationally engineer CoAT to enhance control over MCCA chain length. We anticipate that achieving these interventions could be accelerated by bottom-up microbiome engineering^80^, enabling more precise MCCA upcycling for a sustainable bioeconomy.

## Methods

### Bacterial strains and culturing conditions

*Pseudoramibacter alactolyticus* DSM 3980 and *Anaerostipes caccae* DSM 14662 were purchased from the German Collection of Micro-organisms and Cell Cultures (DSMZ). Both organisms were cultured in Modified Reinforced Clostridial Medium (ATCC 2107) with minor modifications. Starch was removed and glucose was replaced for lactic acid. Chemicals were purchased from BioShop (Burlington, ON, Canada) unless otherwise stated. Beef extract, 10 g L^−1^; Tryptose (Milipore, Burlington, MA, USA), 10 g L^−1^; NaCl, 5 g L^−1^; yeast extract, 3 g L^−1^; sodium acetate, varied concentrations; lactic acid, varied concentrations; resazurin, 0.01% (Sigma-Aldrich, St. Louis, MO, USA); L-cysteine.HCl, 0.5 gL^−1^. pH was adjusted to 6.8. After autoclaving, culture medium was purged with N_2_/CO_2_ gas mix (80%/20%). Batch kinetics were conducted in media bottles sealed with rubber stoppers. Acetate-to-lactate ratio experiment was conducted in 96-well plates. 1-^13^C acetate tracing and cell growth for proteomics were conducted in Hungate tubes. Both strains were statically incubated at 37°C in all experiments.

### Lactate, formate and carboxylic acids quantification

All samples were filtered through a 0.2 μm membrane and stored at −20 °C. Prior to analysis, samples were thawed, centrifuged at 17,000 × g for 2 minutes, and diluted appropriately with HPLC-grade water. Lactate and formate were quantified using high-performance liquid chromatography (HPLC) on an UltiMate 3000 system (Thermo Fisher Scientific, Waltham, MA, USA) equipped with an Aminex HPX-87H column (300 × 7.8 mm, 9 μm, Bio-Rad, Hercules, CA, USA) and a compatible guard column. The mobile phase consisted of 5 mM sulfuric acid, and elution was performed at a flow rate of 0.6 mL/min. The column was maintained at 50 °C. Ultraviolet (UV) detection was used for compound identification and quantification. Data acquisition and analysis were carried out using Chromeleon 7 software (Thermo).

Acetate, butyrate, hexanoate and octanoate were measured using gas chromatography (8900 GC System, Agilent, Santa Clara, CA, USA) with tandem mass spectrometry (7000D Triple Quadrupole GC-MS, Agilent) and DB-FatWax column (Agilent). Carboxylic acids in the standards and diluted samples were protonated with formic acid to a final concentration of 0.1 M. 0.3 µL of each sample were injected with a split ratio of 10:1. The run time was 16 min and the temperature gradient as it follows: 80°C for 1 minute, a 20°C/min ramp until 200°C, 5°C/min until 210°C, 20°C/min until 250°C and finally held for 5 mins. The carrier gas was helium at a flow rate of 1 mL/min. The mass spectrometer was operated either in scan mode (96-well batches and acetate tracing) or dynamic multiple reaction monitoring (dMRM) mode (Batch kinetics). Target analytes were ionized and fragmented by electron ionization. The precursor ion and product ion of target analytes (dMRM) were selected using Agilent software (MassHunter Optimizer, Agilent). To quantify ^13^C labeled acetate, m/z 61 was used while m/z 60 was used for unlabelled acetate.

### Protein extraction and digestion

Biomass was collected during mid-exponential phase by centrifugation, washed once with Phosphate Buffered Saline (PBS), pH 7, and frozen with liquid N_2_. Cell pellets were stored at −80°C until further processing. Protein was extracted and digested as described by Lawson et al.^81^,with minor modifications. Cell pellets were resuspended in 175 µL of 1% sodium deoxycholate in 50 mM triethylammonium bicarbonate (TEAB) buffer, pH 8, and mixed 1:1 (v:v) with B-PER (Thermo Fisher). Cells were lysed with 100 µm acid washed glass beads (Sigma) by vortexing at 2500 rpm for 1 minute. This process was repeated 4 more times with intermittent incubation on ice for 30 seconds. Samples were then frozen at −80°C to aid the lysis process and slowly thawed on ice. Once thawed, samples were vortexed for 1 minute and centrifuged at 14000 rpm for 10 minutes at 4°C. Protein quantification was carried out with Pierce™ Bradford Protein Assay Kit (Thermo Fisher). Cell lysate was then transferred to new tubes and proteins were normalized by mass for differential proteomics. Next, proteins were precipitated with trichloroacetic acid (TCA) at a final concentration of 20% and incubated on ice for 10 minutes. Samples were then centrifuged at 4°C and 14000 rpm for 5 minutes and protein pellet was washed twice with ice cold acetone. Protein pellets were resuspended in 50 µL of 6M urea in 200 mM ammonium bicarbonate (ABC) buffer. Next, dithiothreitol (DTT) in 200 mM ABC buffer was added to a final concentration of 2.3 mM and samples were incubated at 37°C for 1 hour. Next, iodoacetamide (IAM) in 200 mM ABC was added to a final concentration of 3.75 mM and samples were incubated at room temperature, in the dark, for 30 minutes. Prior to tryptic digest, samples were diluted with 200 mM ABC to reach a urea concentration below 1M. Sequencing-grade trypsin (Promega, Madison, WI, USA), 1:50 ratio trypsin to protein, was then added and samples were incubated overnight at 37°C and 350 rpm. Peptide purification was carried out with OMIX Tips (Agilent). Trifluoroacetic acid was added to the samples to a final concentration of 1%. Next, OMIX C18 Tip was wet twice with 100 µL of a 1:1 mixture of acetonitrile (ACN) and H_2_O. Tip was equilibrated twice with 100 µL of 0.1% TFA and samples were aspirated and dispensed 3-5 times. Next, tip was washed twice with 100 µL of 0.1% TFA. Samples were eluted by aspiring and dispensing 100 µL of 95% ACN/0.1% formic acid. Finally, samples were dried in speedvac and resuspended in 100 µL of 0.1% formic acid and filtered with 0.2 µm prior to injection.

### LC/MS parameters for peptide separation and detection

All peptide samples were analyzed using a one-dimensional nano-flow LC-MS/MS setup. Briefly, 10 µL of each sample was injected into an EASY-nano LC 1000 system coupled to a Q-Exactive Orbitrap mass spectrometer (Thermo Scientific) via a nano-electrospray ionization (nanoESI) source. Peptides were separated on a custom-packed SilicaTip emitter column (New Objective, Woburn, USA) using ReproSil-Pur C18-AQ resin (Dr. Maisch GmbH, Ammerbuch-Entringen, Baden-Württemberg, Germany). The mobile phases consisted of solvent A (0.1% formic acid in H_2_O) and solvent B (0.1% formic acid in acetonitrile), and separation was achieved at a constant flow rate of 250 nL/min. A gradient was applied as follows: 0 to 5 min 0% B, increase to 10% B; 5 to 93 min, increase to 40% B; 93 to 95 min, ramp to 95% B; held at 95% B until 105 min; returned to 0% B by 106 min and held until 120 min for column re-equilibration. The mass spectrometer operated in positive ion mode with a spray voltage of 3 kV and a capillary temperature of 275 °C. Full MS scans were acquired from 400 to 2,000 m/z at a resolution of 70,000, with an AGC target of 1e6 and a maximum injection time of 30 ms. For MS/MS acquisition, a Top10 data-dependent method was employed, isolating the top 10 most intense precursor ions within a 0.4 m/z isolation window. MS2 scans were acquired from 200 to 2,000 m/z at a resolution of 17,500, with a maximum injection time of 50 ms and a normalized collision energy (NCE) of 27.

### Raw mass spectra data processing

Tandem mass spectra were extracted and converted to peak lists using the Global Proteome Machine (GPM; thegpm.org) pipeline (version 2020.11.12.1). Charge state deconvolution and deisotoping were not performed. MS/MS spectra were searched using X! Tandem (version X! Tandem Aspartate 2020.11.12.1) against custom protein databases for *P. alactolyticus* (NCBI TaxID: 113287) and *A. caccae* (NCBI TaxID: 105841), containing 2,285 and 3,294 protein entries, respectively. Protein FASTA files were obtained from the NCBI RefSeq database using genome assemblies ASM18550v1 (downloaded in February 2023) and ASM2018143v1 (downloaded in November 2023). Database searches were performed using trypsin as the digestion enzyme, with a precursor ion mass tolerance of 20 ppm and a fragment ion mass tolerance of 0.40 Da. Carbamidomethylation of cysteine and selenocysteine residues was specified as a fixed modification. Variable modifications included N-terminal pyro-glutamate formation from glutamine and glutamic acid, N-terminal ammonia-loss, deamidation of asparagine and glutamine, and oxidation or dioxidation of methionine and tryptophan residues.

Scaffold (version Scaffold_5.1.2, Proteome Software Inc., Portland, OR) was used to validate MS/MS based peptide and protein identifications. Peptide identifications were accepted if they could be established at greater than 95.0% probability by the Peptide Prophet algorithm^82^ with Scaffold delta-mass correction. Protein identifications were accepted if they could be established at greater than 99.0% probability and contained at least 2 identified peptides. Protein probabilities were assigned by the Protein Prophet algorithm^83^. Proteins that contained similar peptides and could not be differentiated based on MS/MS analysis alone were grouped to satisfy the principles of parsimony. Up or downregulation of proteins were assessed using Top 3 total ion current (TIC) as quantitative metric. The t-test was corrected with Benjamini-Hochberg method.

### Cell lysate colorimetric assay

*P. alactolyticus*’ and *A. caccae*’s cell pellets were collected at mid exponential phase, normalized by OD600 (0.8), washed once with 1x PBS, pH7 and stored at −80°C until further processing. 10 mM Tris-HCl buffer (pH 7.5) containing 1 mM DTT and 40 mg/ml of lysozyme were added and samples were incubated at 37 °C for 20 minutes. Next, 0.1g of 100 µm acid washed glass beads (Sigma) were added to aid cell lysis and samples were vortexed at 2500 rpm for 1 minute, followed by ice incubation for 30 seconds. Vortex and ice incubation cycles were repeated 5 more times. Lysate was centrifuged at 13500 rpm for 30 minutes at 4°C and clarified lysate was ultrafiltrated with a 30 kDa MWCO concentrator for 30 minutes at 4°C. CoAT assay was coupled to citrate synthase assay to detect free CoA, as described by Sato et al.^84^, with minor modifications. Briefly, 26 µL of lysate was combined with 10 µL of 40 mM phosphate buffer, pH 7.4, 2 µL of 10 mM acyl-CoA (Sigma) and 2 µL of 2M sodium acetate. Samples were incubated at 25°C for 5 min. Next, 40 µL of 8 mM oxaloacetic acid, 8 mM of 5,5′-dithiobis-2-nitrobenzoic acid (DTNB) and 46.5 μg/mL of citrate synthase (NZYTech, Lisbon, Portugal) in 400mM phosphate buffer, pH 8, was added to each assay. Samples were incubated at 37°C for 30 min. Absorbance was read at 412 nm and negative controls for each condition did not have sodium acetate added. Acetyl-CoA formation quantification was evaluated using acetyl-CoA (Sigma) to prepare a calibration curve.

### Enzyme Cost Minimization (ECM)

Python packages equilibrator-api and equilibrator-pathway (https://gitlab.com/equilibrator) were used to perform ECM, developed by Flamholz *et al*^43^. A reaction’s enzyme demand [*E*] (molar) given flux *J* is:

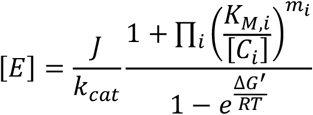

*k*_*cat*_ is the forward rate constant, [*C*_*i*_] is the concentration of reactant i, *K*_*M*,*i*_ is the enzyme’s saturation constant for reactant i, *m*_*i*_ is the stoichiometric coefficient of reactant i, and Δ*G*^′^ is the Gibbs free energy change, estimated via component contribution at pH 6^85^. Total enzyme cost of a pathway per unit flux is minimized:

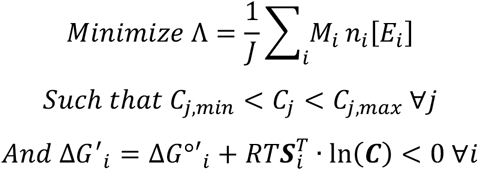

Where ***S*** is the pathway’s stoichiometric matrix and ***C*** contains metabolite concentrations. For the i^th^ reaction, *n*_*i*_ is its relative flux, *M*_*i*_ is its enzyme’s molar weight. Rate constants and saturation constants are shared across all enzymes (k_cat_ = 200 s^-1^; K_m_ = 0.2 μM), molecular weights (*MW*) were taken from Uniprot (https://www.uniprot.org/). Lower and upper bounds of pathway specific metabolites are 1 μM and 10 mM, apart from oxoacyl-CoAs (lower bound of 0.4 μM and upper bound of 10 mM). Universal metabolite (e.g. ATP) concentrations are derived from literature^86^. The reduction potential of ferredoxin is set to −400 mV and NADH to −280 mV. Saturation effects of redox equivalents are omitted. RNF, HYDA and ATP synthase were not included in enzyme cost estimates.

### Stoichiometric Model

*n* is the mean number of RBO cycles, derived from measurements of butyrate ([*B*]), hexanoate ([*H*]) and octanoate ([*O*]) production:

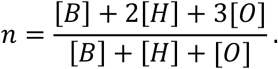

*θ* is the SLP flux per product flux (specific flux); *θ* = 0 corresponds to complete acetate assimilation and *θ* = 1 to complete recycling. Molar product yield on acetate 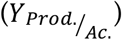 is related to acetate recycling through:

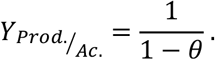

*θ* can therefore be estimated from butyrate, hexanoate and octanoate production and acetate consumption data.

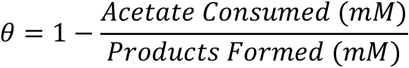

Specific lactate consumption flux is *n* + *θ*: increased acetate recycling requires that more lactate is oxidized to supplement acetyl-CoA lost to SLP. Specific RNF flux, *α*, is the difference of NADH production via lactate oxidation to acetyl-CoA and NADH consumption by RBO cycles per mole of product. Since reverse flux through RNF is predicted to be infeasible:

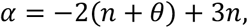

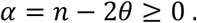

Values of *n* and *θ* that satisfy this inequality correspond to combinations of product profile and acetate utilization strategy for which NADH balances; if the inequality is violated, redox balance would require negative flux through RNF to balance a surplus of NADH.

If a fraction of RBO cycles (*ε*) use an NADPH-dependent HACD and NFN to regenerate NADPH from NADH and reduced ferredoxin via electron confurcation, specific NFN flux equals *nε* and specific RNF flux becomes:

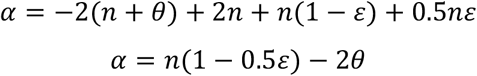

This reduces to the previous expression when *ε* = 0.

Specific HYDA flux is equal to ferredoxin reduced by EB-ACD minus ferredoxin oxidized by RNF and NFN, and ferredoxin diverted by PFL per mole of product:

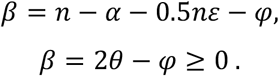

ATP yield per mole of lactate is one half RNF flux plus SLP flux per lactate flux:

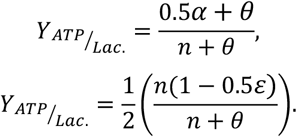

And if NFN flux is negligeable:

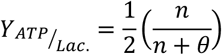

See Supplementary Material S1 for calculations.

## Data availability

The mass spectrometry proteomics data have been deposited to the ProteomeXchange Consortium via the PRIDE^87^ partner repository with the dataset identifier PXD063079 and 10.6019/PXD063079. ECM models and python functions are available on GitHub (https://github.com/Connor-Bowers/Acetate-Utilization-Strategy).

## Acknowledgements

The authors would like to acknowledge Joel Howard, Jasmeen Parmar and Diana Dyussekenova for the support on equipment operation. This work was supported by the Natural Sciences and Engineering Research Council of Canada [ALLRP 580897-22, RGPIN-2021-02684, NSERC-CREATE 528163-2019].

