## Supplementary Material for "Distinct Acetate Utilization Strategies Differentiate Butyrate and Medium-Chain Carboxylic Acid Producing Chain-Elongating Bacteria"

**Supplementary Text**

*ATP yield predictions*

Acetate recycling has important implications for energy conservation in RBO. Chemiosmosis is often assumed to be the primary mode of energy conservation in CEB, with SLP playing a minor and static role. These results show that SLP becomes increasingly prevalent in both *A. caccae* and *P. alactolyticus* as acetate supplementation is reduced. Notably, stoichiometric analysis suggests that acetate recycling reduces ATP yield on lactate by decreasing chemiosmosis through RNF ($\alpha$) and increasing lactate utilization. As stated in the Methods, RNF flux ($\alpha$) decreases with acetate recycling according to:

$$\alpha=n-2\theta$$

Where $n$ is the mean number of RBO cycles, and $\theta$ is specific flux of acetate recycling (moles acetate recycled per mole of product).

ATP yield per mole of lactate is then one half RNF flux ($0.5\alpha$) plus SLP flux ($\theta$) divided by lactate flux ($n+\theta$). Therefore, we see that ATP yield decreases inversely with acetate recycling according to:

$$Y_{\frac{ATP}{Lac.}}=\frac{0.5\alpha+\theta}{n+\theta},$$

$$Y_{\frac{ATP}{Lac.}}=\frac{1}{2}\left( \frac{n}{n+\theta} \right).$$

On rich media, *A. caccae* favoured acetate assimilation over recycling and its predicted RNF flux and ATP yield tended to be higher than *P. alactolyticus* (Fig. 1f, Supplementary Material Fig. 1b). Interestingly, when grown on dilute media, *A. caccae* recycled more acetate relative to rich media, while *P. alactolyticus* assimilated more relative to rich media. As a result, predicted ATP yield and RNF flux were similar for both organisms (Supplementary Material Fig. 1c and Fig. 9f). This implies that under nutrient limited conditions, *A. caccae* does not strictly maximize growth yield, and instead recycles some acetate to maintain lactate uptake as acetate is depleted. Importantly though, *A. caccae* was still unable to recycle more than 0.5 moles of acetate per mole of product ($\theta=0.5$) and did not grow at acetate-to-lactate ratios below 0.1. On the other hand, *P. alactolyticus* maintained growth in these conditions by fully recycling acetate ($\theta=1$). Taken together, these results suggest that acetate recycling imposes an ATP yield penalty and can require MCCA production ($\theta>0.5$). Under acetate limitation, increased lactate utilization enabled by strong recycling may outweigh these potential drawbacks.

**Supplementary Material S1**

Calculation of reverse beta-oxidation (RBO) fluxes per product flux (lactate oxidation, RBO cycle reactions, substrate level phosphorylation, proton pumping) from product formation and acetate consumption data for *P. alactolyticus* and *A. caccae* grown on varied ratios of acetate and lactate. Calculation of hydrogen and formate evolution fluxes per product flux from formate and product formation and acetate consumption at select acetate-to-lactate ratios.

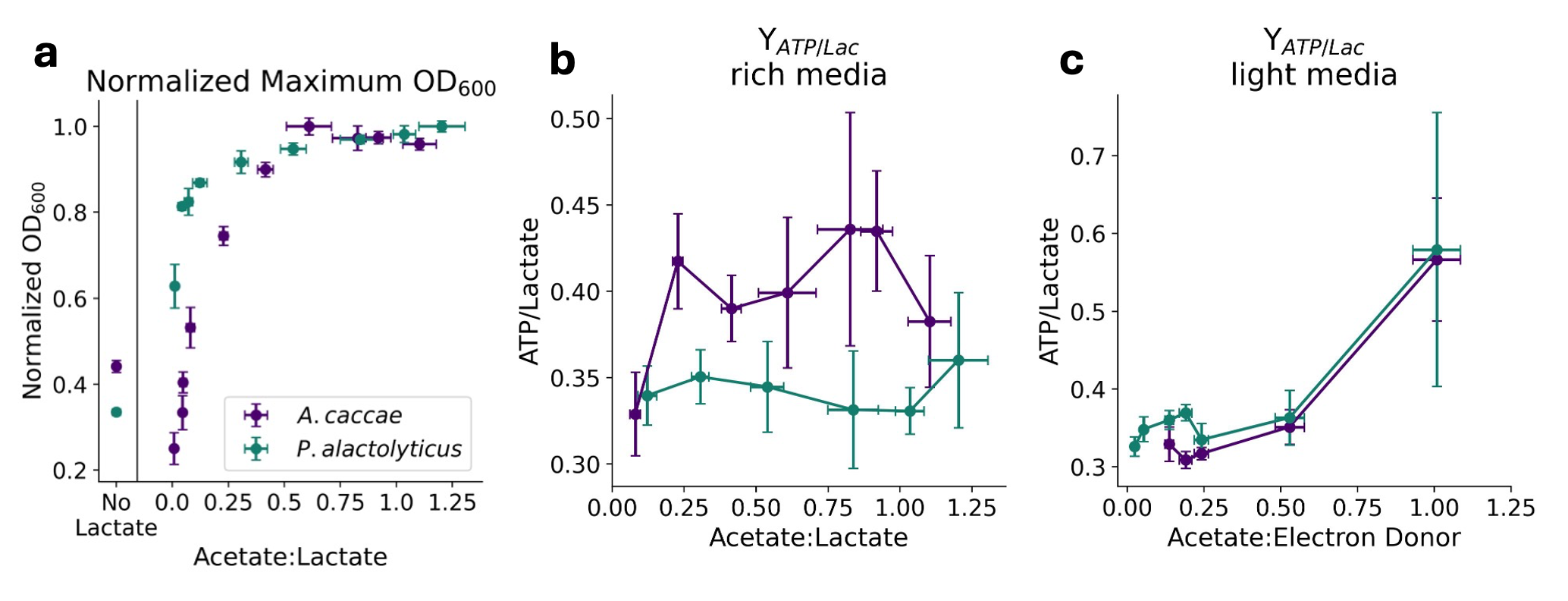

**Supplementary Fig. 1.|** **a.** Maximum optical density (OD_600_) achieved by *P. alactolyticus* and *A. caccae* when grown on varied ratios of acetate and lactate, keeping lactate constant. Maximum reading for each ratio is normalized to the maximum across ratios. Values compared to OD_600_ without lactate or acetate supplementation. *P. alactolyticus* maintains growth when acetate is limiting better than *A. caccae* by recycling acetate and maximizing lactate utilization. **b.** ATP yield on lactate predicted by stoichiometric model for *P. alactolyticus* and *A. caccae* across acetate-to-lactate ratios. *A. caccae* may extract more ATP by favouring acetate recycling. **c.** ATP yield on lactate predicted with data from dilute media, showing similar values for both organisms, since *A. caccae* displayed more acetate recycling than in rich media (still less than 0.5 moles acetate recycled per mole of product formed), and acetate assimilation by *P. alactolytics* in high acetate conditions was higher than previously observed. All replicates in this figure are biological triplicates. Error bars represent 1 standard deviation.

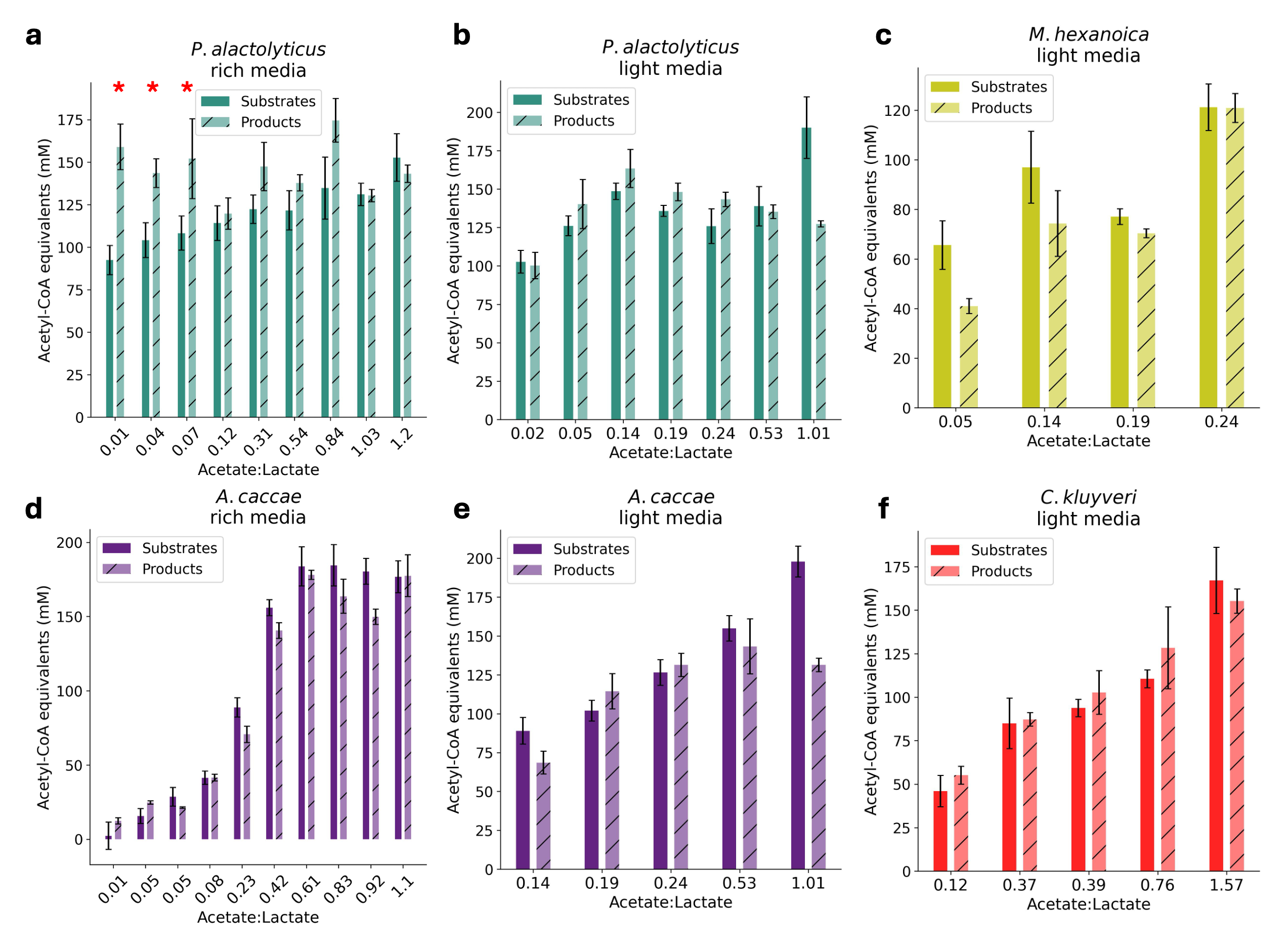

**Supplementary Fig. 2.|Acetyl-CoA equivalents (AcCoA eq) in substrates (acetate and lactate) and products (butyrate, hexanoate and octanoate) across acetate to lactate ratios. a.** *P. alactolyticus* balance on rich ATCC 2107. AcCoA eq in products were significantly higher than in lactate and acetate consumed for the three lowest acetate-to-lactate ratios (0.01, 0.04, 0.07), suggesting consumption of an unidentified carbon source.  **b.** *P. alactolyticus* balance on ATCC 2107 Lite **c.** *M. hexanoica* balance on ATCC 2107 Lite **d.** *A. caccae* balance on rich ATCC 2107 **e.** *A. caccae* balance on ATCC 2107 Lite **f.** *C. kluyveri* balance on ATCC 2107 Lite.

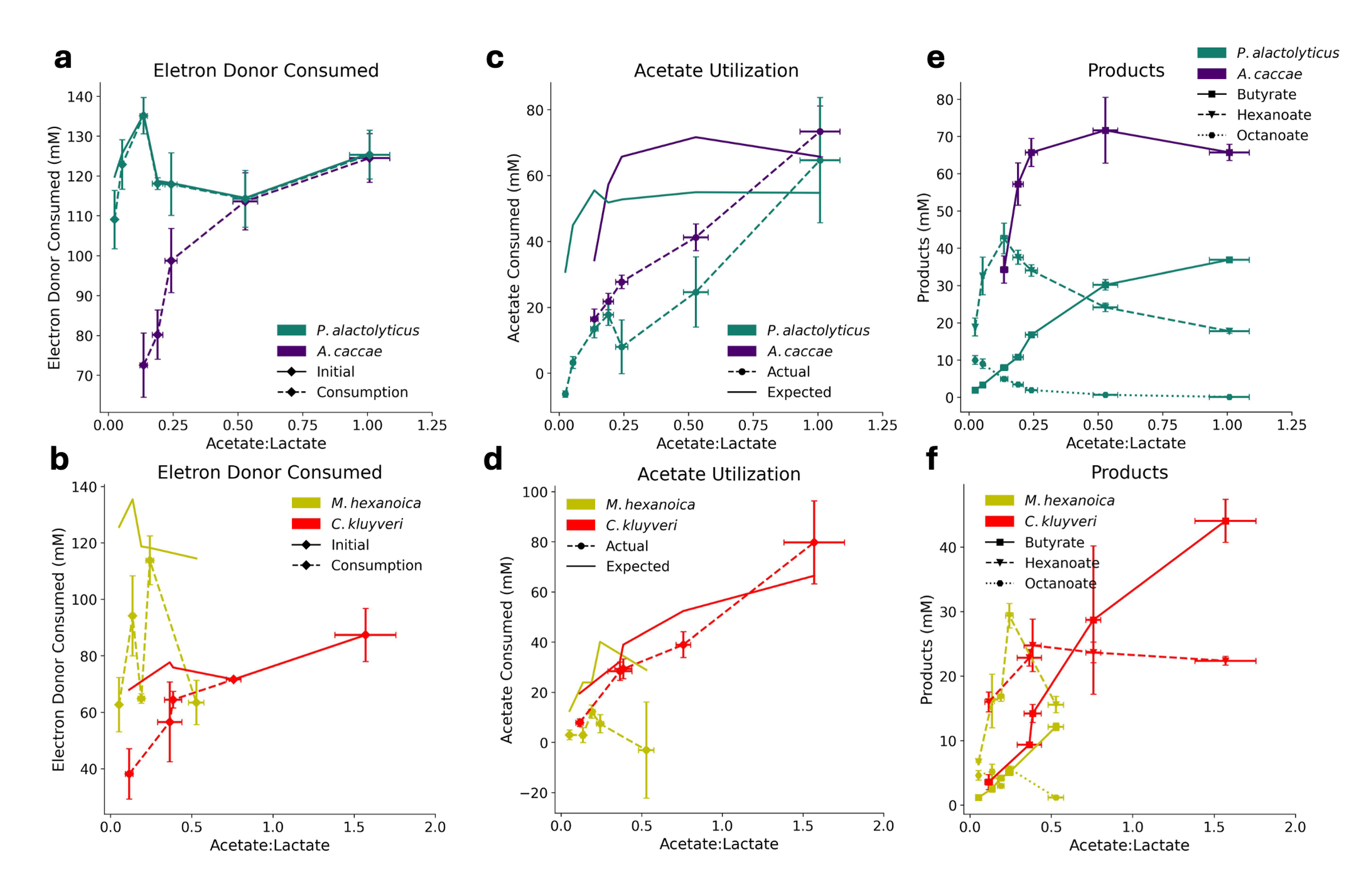

**Supplementary Fig. 3.|Electron donor utilization, acetate utilization and product spectra in four CEB in ATCC 2107 Lite under various acetate-to-electron donor ratios**. **a.** Initial lactate concentration and lactate utilization by *P. alactolyticus* and *A. caccae*. *P. alactolyticus* maintains lactate utilization across conditions thanks to acetate recycling. **b.** Initial eletron donor concentration and electron donor utilization by *M. hexanoica* (lactate) and *C. kluyveri* (ethanol). **c.** Expected and actual acetate consumption by *P. alactolyticus*, *A. caccae* and **d.** *M. hexanoica* and *C. kluyveri*.  **e.** Carboxylic acid production by *P. alactolyticus*, *A. caccae* and **f.** *M. hexanoica* and *C. kluyveri*.

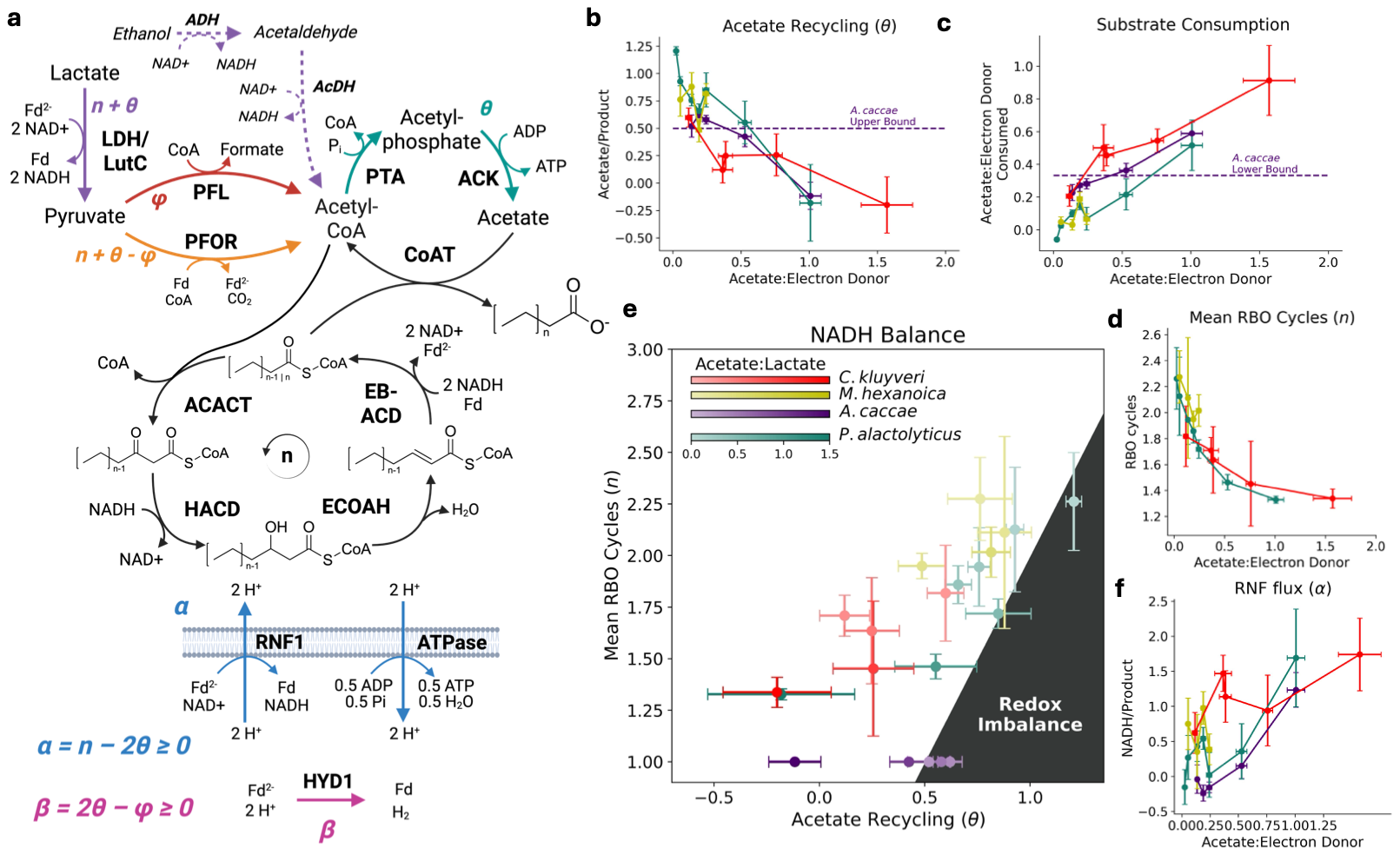

**Supplementary Fig. 4.|Extending stoichiometry modelling to more CEB in ATCC 2107 Lite media corroborates the relationship between strong acetate recycling and elongation to MCCAs. a.** Stoichiometric model of RBO with simultaneous acetate assimilation and recycling, where $\theta$ is the extent of acetate recycling (moles recycled per mole product). LDH = electron bifurcating lactate dehydrogenase; LutC = lactate utilization protein C; PFOR = pyruvate:ferredoxin oxidoreductase; ADH= alcohol dehydrogenase; AcDH= acetaldehyde dehydrogenase; PFL = pyruvate formate lyase; PTA = phosphate acetyltransferase; ACK = acetate kinase; ACACT = acetyl-CoA C-acetyltransferase; HACD = 3-hydroxyacyl-CoA dehydrogenase; ECOAH = enoyl-CoA hydratase; EB-ACD = electron bifurcating acyl-CoA dehydrogenase, comprising acyl-CoA dehydrogenase (ACD) and electron transfer flavoproteins A and B (EtfA and EtfB); CoAT = acyl-CoA:acetate CoA transferase; Fd = ferredoxin; RNF1 = proton translocating ferredoxin: NAD+ oxidoreductase complex; ATPase: F0/F1 ATP synthase; HYD1 = Fe–Fe hydrogenase. **b.** Acetate recycling as a function of acetate-to-lactate ratio **c.** and ratio of acetate-to-lactate consumption at different ratios of supplemented acetate and lactate. **d.** Mean number of RBO cycles ($n$) in *P. alactolyticus*, *M. hexanoica* and *C. kluyveri* as a function of acetate-to-lactate ratio. **e.** Experimentally derived pairs of $n$ and $\theta$, and inequality relating these parameters for maintenance of redox balance ($\alpha$ > 0). **f.** Flux through RNF relative to product flux ($\alpha$) for *A. caccae*, *P. alactolyticus*, *M. hexanoica* and *C. kluyveri*

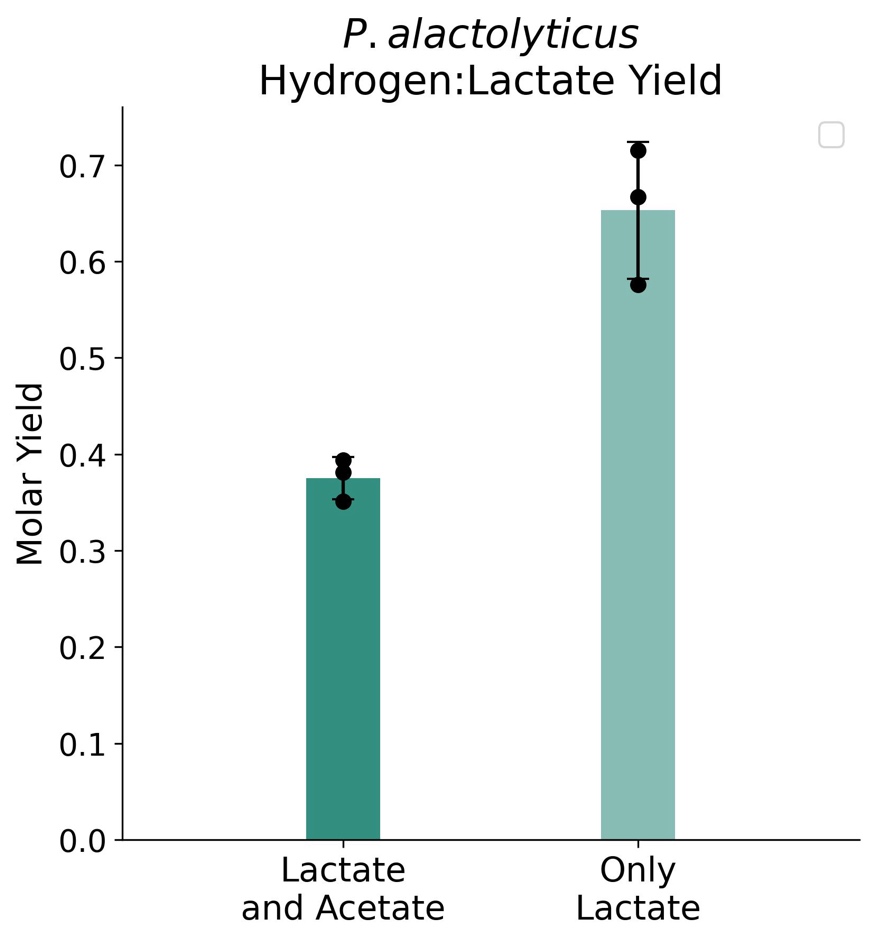

**Supplementary Fig. 5|Hydrogen production increases with acetate recycling.** Hydrogen production from lactate with and without acetate supplementation (n=3). Difference between conditions was validated using a two-tailed Student’s t-tests (p=0.003). All replicates in this figure are biological. Error bars represent 1 standard deviation.

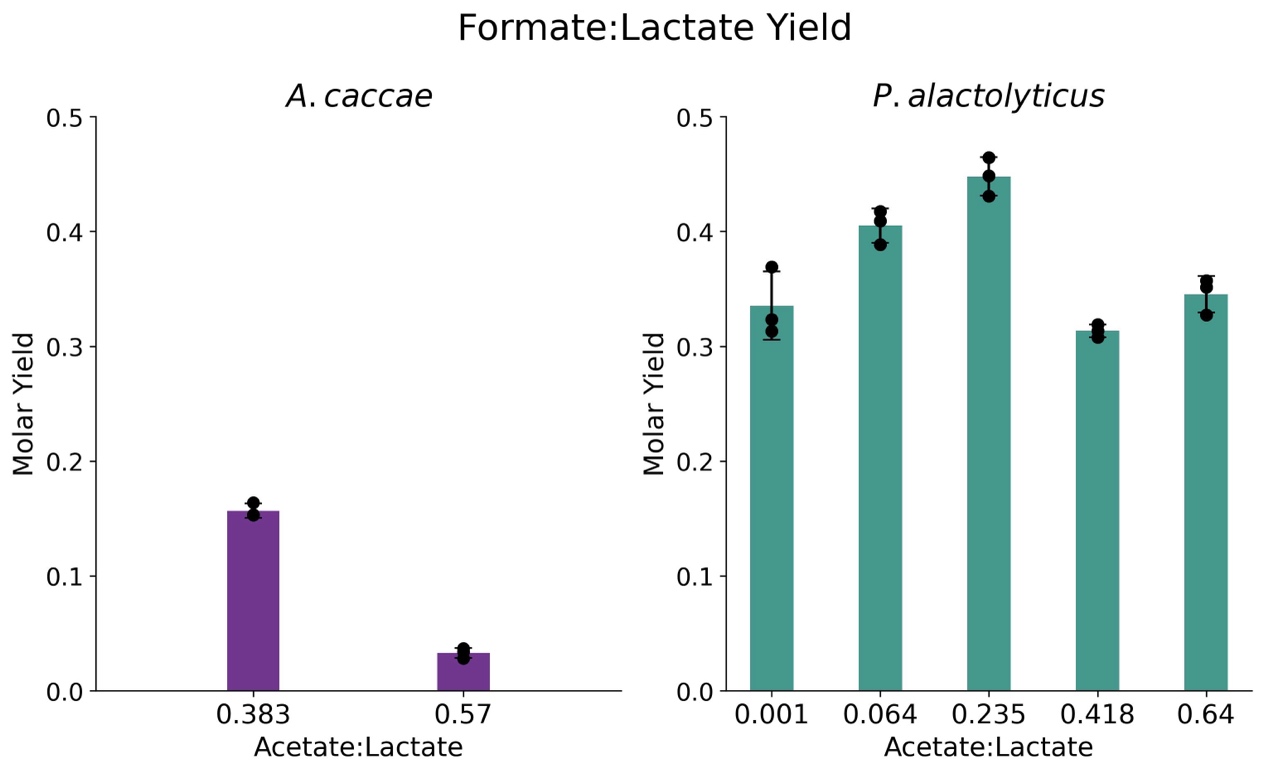

**Supplementary Fig. 6| *P. alactolyticus* has higher PFL flux than *A. caccae* to balance excess of reduced ferredoxin and prevent rapid accumulation of H_2_.** Left: Formate production in *A. caccae* under different acetate-to-lactate ratios. Right: Formate production in *P. alactolyticus* under different acetate-to-lactate ratios. Acetate-to-lactate ratios were determined experimentally. All conditions were performed in triplicate (n=3). All replicates in this figure are biological. Error bars represent 1 standard deviation.
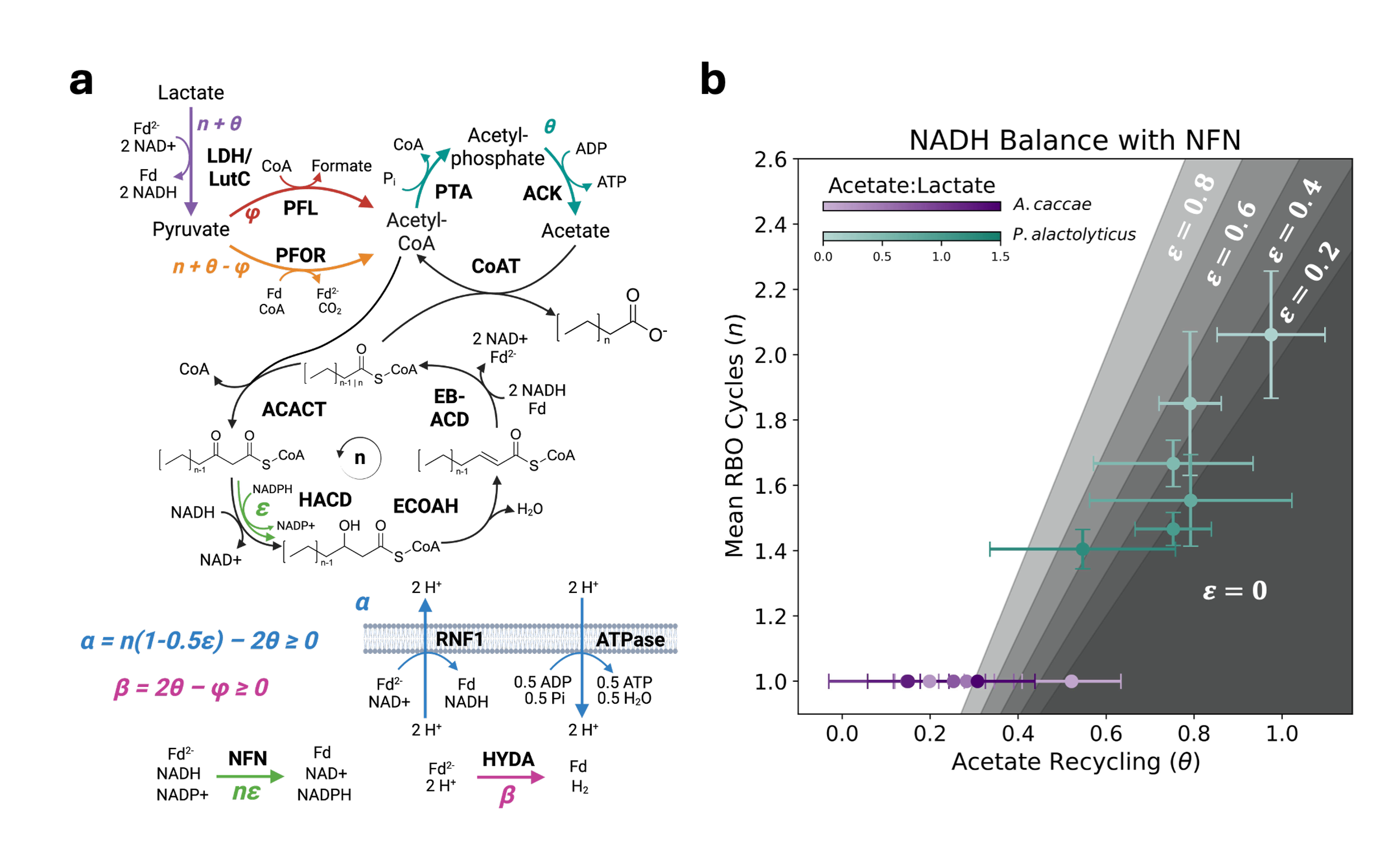

**Supplementary Fig. 7.|Incorporating NADPH dependent 3-hydroxyacyl-CoA dehydrogenase (HACD) and NFN into stoichiometric model strengthens requirement for chain elongation when acetate is recycled. a.** Stoichiometric model of RBO with simultaneous acetate assimilation and recycling. $\varepsilon$ is the fraction of RBO cycles that use an NADPH dependent HACD, and $n\varepsilon$is flux through NFN per product flux. LDH = electron bifurcating lactate dehydrogenase; LutC = lactate utilization protein C; PFOR = pyruvate:ferredoxin oxidoreductase; ADH= alcohol dehydrogenase; AcDH= acetaldehyde dehydrogenase; PFL = pyruvate formate lyase; PTA = phosphate acetyltransferase; ACK = acetate kinase; ACACT = acetyl-CoA C-acetyltransferase; HACD = 3-hydroxyacyl-CoA dehydrogenase; ECOAH = enoyl-CoA hydratase; EB-ACD = electron bifurcating acyl-CoA dehydrogenase, comprising acyl-CoA dehydrogenase (ACD) and electron transfer flavoproteins A and B (EtfA and EtfB); CoAT = acyl-CoA:acetate CoA transferase; Fd = ferredoxin; RNF1 = proton translocating ferredoxin: NAD+ oxidoreductase complex; ATPase: F0/F1 ATP synthase; HYD1 = Fe–Fe hydrogenase. **b.** Experimentally derived pairs of $n$ and $\theta$ for *P. alactolyticus* and *A. caccae*, and inequalities relating these parameters for maintenance of redox balance ($\alpha$ > 0) for different values of $\varepsilon$. If $\varepsilon$ increases, each RBO cycle consumes less NADH, meaning more RBO cycles are required to balance additional NADH from lactate when acetate is recycled. NFN does not appear to play a major role in RBO for the organisms under study, since they respect the redox constraint predicted if NFN flux is negligeable ($\varepsilon\approx0$).

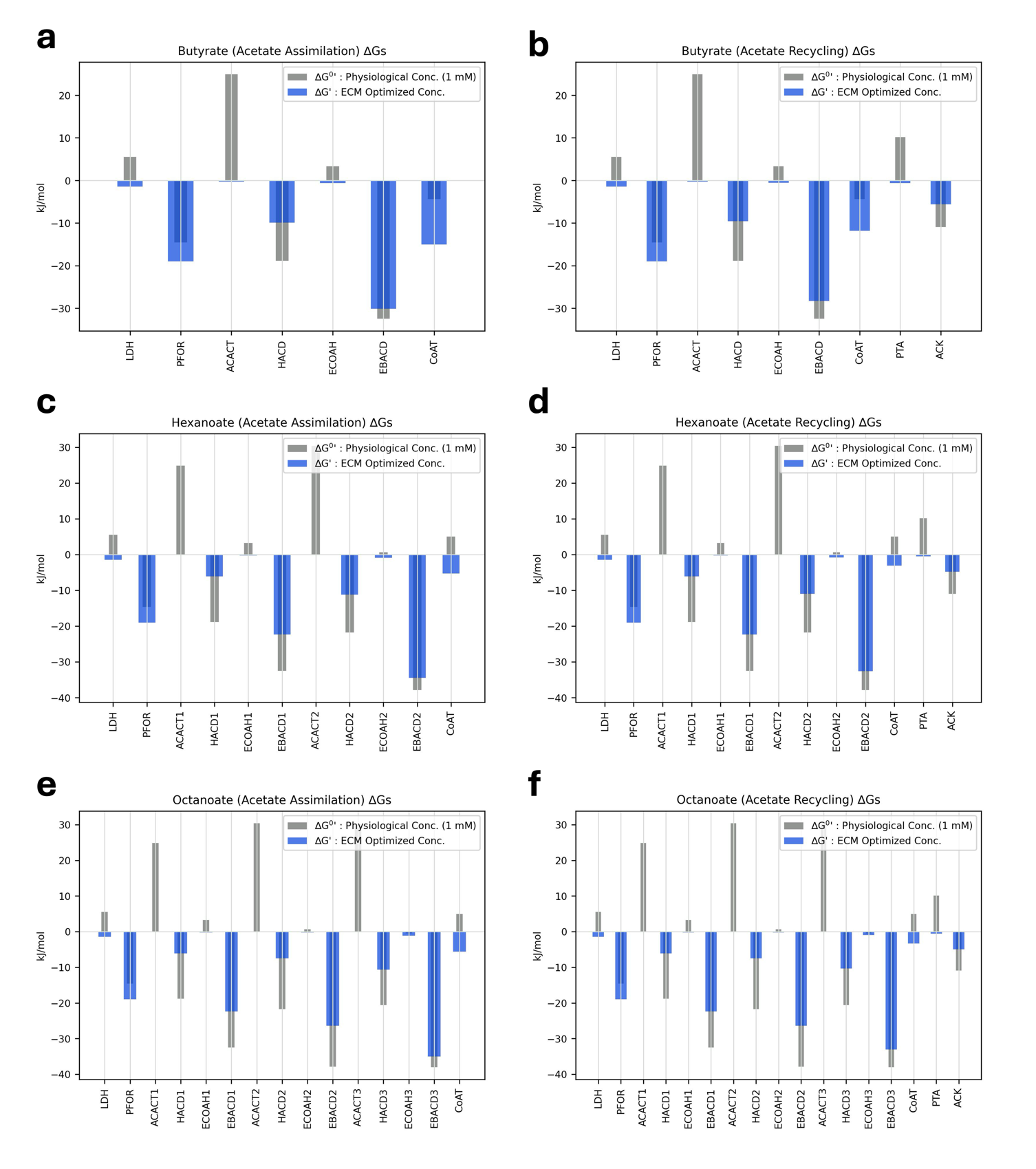

**Supplementary Fig. 8.|** Gibbs free energy change estimates from Enzyme Cost Minimization (ECM) of RBO reactions for butyrate (**a.** and **b.**), hexanoate (**c.** and **d.**) and octanoate (**e.** and **f.**) production via complete acetate assimilation or recycling. Gibbs free energy changes with standard physiological metabolite concentrations (1 mM) are calculated at pH 6 (grey bars). ECM finds the distribution of metabolite concentrations that minimizes total enzyme cost while keeping all reactions exergonic, yielding an estimate of intracellular Gibbs free energy changes (blue bars). ACACT (thiolase) reactions are the thermodynamic bottlenecks of RBO, requiring high substrate concentrations and low product concentrations which reduces the driving force of proceeding (HACD) and preceding (EB-ACD) reactions at the enzyme cost minimum. As a result, RBO reactions become less exergonic as chain length grows and more thiolase reactions are required.

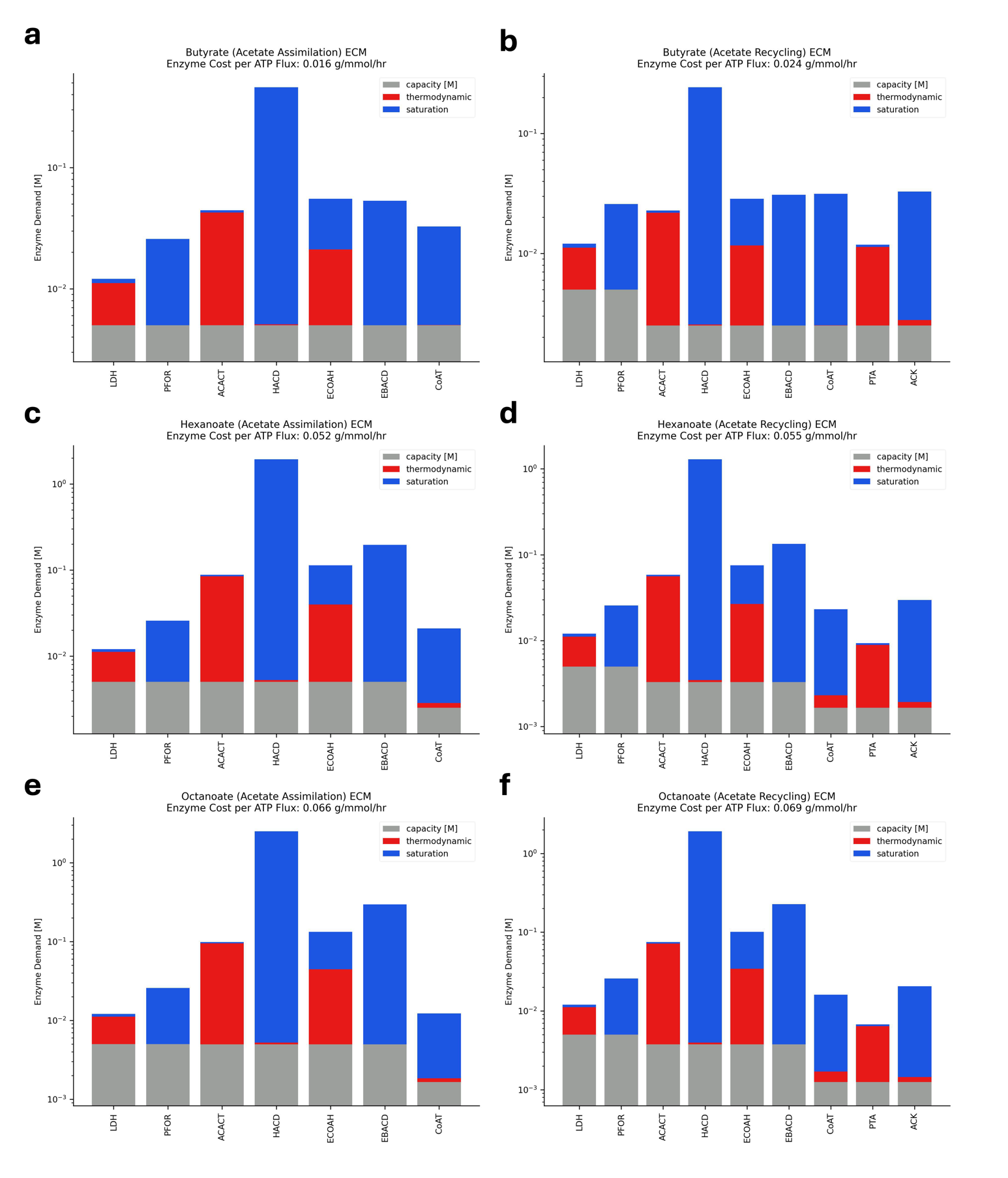

**Supplementary Fig. 9.|** Enzyme Cost Minimization (ECM) of RBO pathways for butyrate (**a.** and **b.**), hexanoate (**c.** and **d.**) and octanoate (**e.** and **f.**) production via complete acetate assimilation or recycling. Cost contributions of stoichiometric effects (grey), proximity to equilibrium (red) and enzyme saturation (blue) are distinguished. Costs of RBO reactions for different cycles (chain lengths) are binned together. Increased number of thiolase reactions and indirect effects of these bottlenecks on neighbouring reactions cause enzyme cost to increase with chain length. PTA and ACK contribute to higher enzyme demand for acetate recycling.

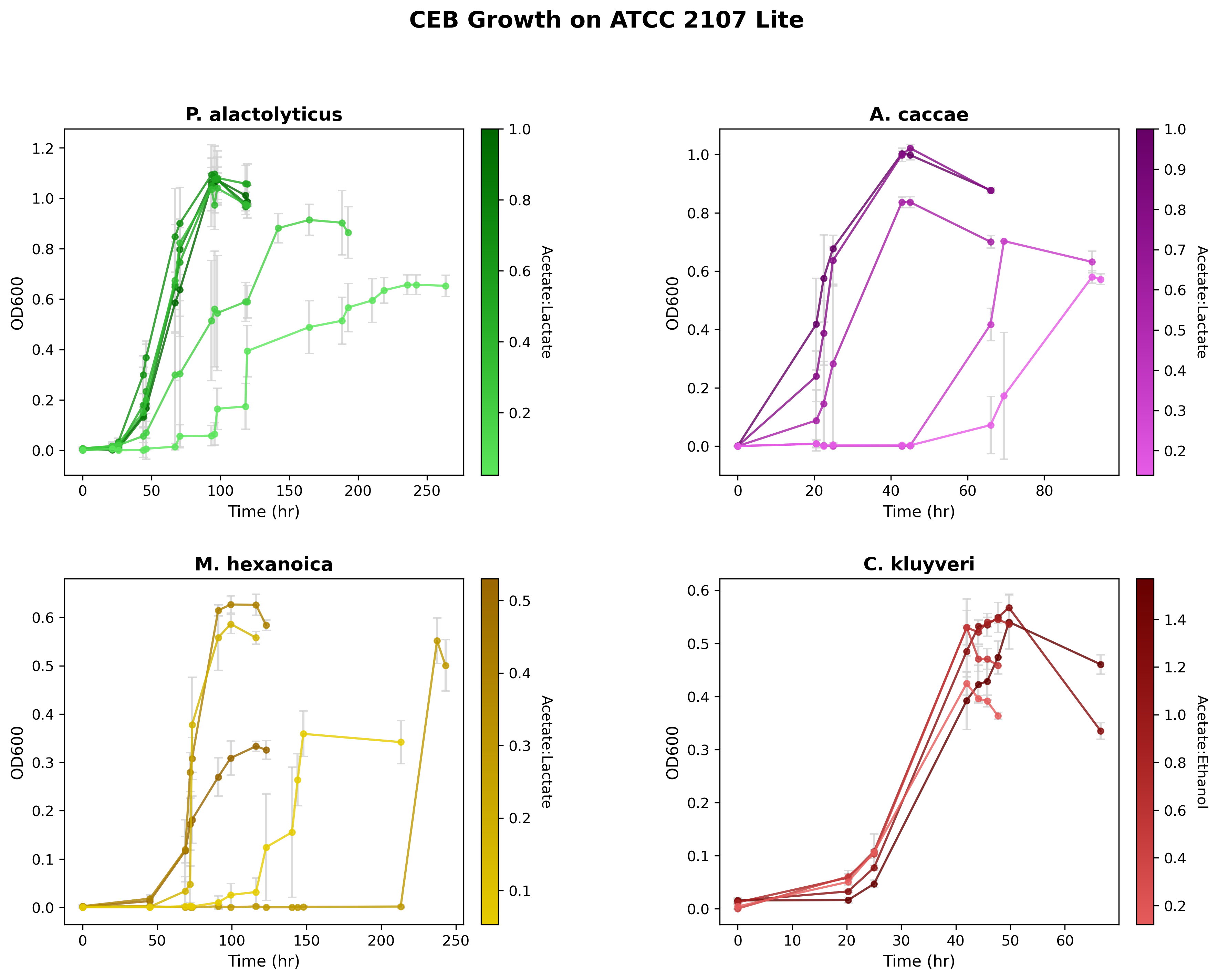

**d**

**c**

**b**

**a**

**Supplementary Fig. 10.| Growth response of four CEB on ATCC 2107 Lite showcases their growth diversity across conditions**. **a.** P. alactolyticus maintained growth under acetate-limiting conditions, exhibiting appreciable growth across all acetate-supplemented treatments. **b.** A. caccae displayed faster growth overall but was unable to grow when acetate concentrations approached ~20 mM. **c.** M. hexanoica showed a growth pattern similar to P. alactolyticus, but exhibited inhibition at high acetate or lactate concentrations and failed to grow when acetate levels were below ~7 mM. **d.** C. kluyveri demonstrated decreasing maximum OD as acetate availability declined and was unable to grow at acetate concentrations below ~8 mM. Error bars represent 1 standard deviation. All data points represent the average of three biological replicates (n=3).

$\theta$ $n$ $n$ $\theta$ $\alpha$ $\alpha$.

**Supplementary Methods**

**Additional CEB strains**

*Clostridium kluyveri* DSM 555 and *Megasphaera hexanoica* DSM 106893 were purchased from the German Collection of Micro-organisms and Cell Cultures (DSMZ).

**Batch fermentations**

Diluted Modified Reinforced Clostridial Medium (ATCC 2107 Lite) consisted of the regular ATCC 2107 used in this manuscript, but with reduced extracts concentrations, as follows. Chemicals were purchased from Bioshop (Burlington, ON, Canada), unless otherwise stated . Beef extract, 1 g L^−1^; Tryptose (Milipore, Burlington, MA, USA), 1 g L^−1^; NaCl, 5 g L^−1^; yeast extract, 1 g L^−1^; sodium acetate, varied concentrations; lactic acid 100 mM; resazurin, 0.01% (Sigma-Aldrich, St. Louis, MO, USA); L-cysteine.HCl, 0.5 gL^−1^. pH was adjusted to 6.8-7.0. To support Clostridium kluyveri growth, lactate was replaced with 100 mM ethanol, and potassium phosphate buffer, 20 mM, p-aminobenzoic acid, 100 µg L^−1^, and biotin, 5 µg L^−1^, were added to the medium. The pH was adjusted to 6.8-7.0.

Batch cultures in rich medium were incubated at 37 °C either statically in Hungate tubes or in borosilicate media bottles, each filled to 50% of the vessel volume. For the 96-well plate experiment, cultures were grown in rich medium inside an Agilent BioTek BioSpa™ live-cell analysis system maintained at 37 °C within an anaerobic chamber. The BioSpa was coupled to an Agilent BioTek plate reader configured for OD₆₀₀ measurements. Prior to each reading, plates were shaken for 1 min using the integrated orbital shaker to homogenize cell suspensions. For the defined-medium ratio experiment, cultures were grown in Hungate tubes due to the substantial medium evaporation observed in microplates after 5 days in the dry anaerobic chamber atmosphere. Tubes were incubated at 37 °C with 80 rpm orbital shaking. *C. kluyveri*’s cultures were incubated statically, as it did not grow when subjected to shaking.

The pH was not controlled for lactate fermentation with A. caccae, P. alactolyticus, and M. hexanoica as the final pH in all batches were within 6.6-7.08 (initial pH ~6.8). Buffer was only added to control pH for ethanol fermentation with C. kluyveri, where the final pH was kept between 5.8-6.0 (initial pH ~6.8-7.0).

**Alternative method for lactate quantification**

Lactate in the ATCC 2107 Lite ratios experiment was quantified using the lactate dehydrogenase assay. Lactate dehydrogenase assays were performed in 96-well plates using lactate standards ranging from 0 to 7 mM (final concentrations). Reaction mixtures contained diluted samples, 0.2 M Bicine buffer (Sigma-Aldrich, St. Louis, MO, USA) and 2.5 mM NAD⁺ (Bioshop, Burlington, ON, Canada). Enzymatic conversion was carried out using 8 U/mL^−1^ of L-lactate dehydrogenase from rabbit muscle (Sigma-Aldrich, St. Louis, MO, USA) and 8 U/mL^−1^ D-lactate dehydrogenase (NZYTech, Lisbon, Portugal). Plates were incubated at 37 °C and monitored kinetically until reactions reached saturation. Absorbance was measured at 340 nm, and the maximum absorbance value obtained for each well was used for calculations.

**Ethanol quantification**

Ethanol in *C. kluyveri* experiments in ATCC 2107 Lite was measured using gas chromatography (8900 GC System, Agilent, Santa Clara, CA, USA) with tandem mass spectrometry (7000D Triple Quadrupole GC-MS, Agilent) and DB-FatWax column (Agilent), operating in headspace mode. Prior to injection, diluted samples and standards were heated at 95°C for 40 minutes in a tightly sealed vial. The sample for injection was taken from the gas phase of the vial. The run time was 12 minutes, with a 1.5 minute solvent delay and the following temperature gradient: 80°C for 1 minute, followed by a 20°C min-1 ramp until 200°C, and held for 5 minutes. The carrier gas was helium at a flow rate of 1 mL/min. The mass spectrometer was operated in scan mode. Target analytes were ionized and fragmented by electron ionization. The precursor ion and product ion of target analytes (dMRM) were selected using Agilent software (MassHunter Optimizer, Agilent).

**Carboxylic acids extraction with ethyl acetate**

Ethyl acetate extractions were required for the experiments using ATCC 2107 Lite. Richness of the original ATCC 2107 likely helped stabilizing C4, C6 and C8 in solution after thawing. This did not happen in ATCC 2107 Lite, as C4-C8 were underestimated using aqueous injections. Samples were diluted, to fall within the GC-MS/MS limits, followed by acidification with HCl to a final concentration of 0.1N. Ethyl acetate was then added at the same volume as the aqueous phase and vortexed for 3 minutes at max speed. Organic and aqueous phase separation was aided by centrifugation at 13000 rpm for 2 minutes. The organic layer was finally transferred to vials for GC-MS/MS quantification. Standards followed the same procedure for acidification and extraction. Butyrate, hexanoate and octanoate were measured using gas chromatography (8900 GC System, Agilent, Santa Clara, CA, USA) with tandem mass spectrometry (7000D Triple Quadrupole GC-MS, Agilent) and DB-FatWax column (Agilent). 0.3 mL of each sample were injected with a split ratio of 10:1. The run time was 20 min and the temperature gradient as it follows: 120°C for 2 minutes, a 5°C/min ramp until 140°C and held for 3 minutes, 20°C/min until 250°C and finally held for 5.5 minutes. The carrier gas w as helium at a flow rate of 1 mL/min. The mass spectrometer was operated in scan mode. Target analytes were ionized and fragmented by electron ionization. The precursor ion and product ion of target analytes (dMRM) were selected using Agilent software (MassHunter Optimizer, Agilent).

**Hydrogen quantification**

Hydrogen partial pressure was measured using a PerkinElmer Clarus 680 gas chromatograph equipped with a thermal conductivity detector (TCD) and a Carboxen 1000 column (Supelco, Bellefonte, USA). Argon was used as the carrier gas at a flow rate of 30 ± 0.1 mL/min. The oven program was 30 °C for 5 min, ramped at 20 °C min⁻¹ to 225 °C. Samples (100 µL) were injected using a gas-tight glass syringe (Chromatographic Specialties, Brockville, Canada). Ideal gas was assumed for quantification.
